# Quantifying the extent to which index event biases influence large genetic association studies

**DOI:** 10.1101/074781

**Authors:** Hanieh Yaghootkar, Michael P. Bancks, Sam E. Jones, Aaron McDaid, Robin Beaumont, Louise Donnelly, Andrew R. Wood, Archie Campbell, Jessica Tyrrell, Lynne J. Hocking, Marcus A. Tuke, Katherine S. Ruth, Ewan R. Pearson, Anna Murray, Rachel M. Freathy, Patricia B. Munroe, Caroline Hayward, Colin Palmer, Michael N. Weedon, James S. Pankow, Timothy M. Frayling, Zoltán Kutalik

## Abstract

As genetic association studies increase in size to 100,000s of individuals, subtle biases may influence conclusions. One possible bias is “index event bias” (IEB), also called “collider bias”, caused by the stratification by, or enrichment for, disease status when testing associations between gene variants and a disease-associated trait. We first provided a statistical framework for quantifying IEB then identified real examples of IEB in a range of study and analytical designs. We observed evidence of biased associations for some disease alleles and genetic risk scores, even in population-based studies. For example, a genetic risk score consisting of type 2 diabetes variants was associated with lower BMI in 113,203 type 2 diabetes controls from the population based UK Biobank study (−0.010 SDs BMI per allele, P=5E-4), entirely driven by IEB. Three of 11 individual type 2 diabetes risk alleles, and 10 of 25 hypertension alleles were associated with lower BMI at p<0.05 in UK Biobank when analyzing disease free individuals only, of which six hypertension alleles remained associated at p<0.05 after correction for IEB. Our analysis suggested that the associations between *CCND2* and *TCF7L2* diabetes risk alleles and BMI could (at least partially) be explained by IEB. Variants remaining associated after correction may be pleiotropic and include those in *CYP17A1* (allele associated with hypertension risk and lower BMI). In conclusion, IEB may result in false positive or negative associations in very large studies stratified or strongly enriched for/against disease cases.

## Introduction

Genome wide association studies (GWAS) in increasingly large sample sizes have resulted in the identification of many 100s of genetic variants associated with common diseases^1–3^. We assume that the results from GWAS are robust to potential confounding factors and biases because genetic variants are inherited randomly and not influenced by disease processes throughout life. These assumptions tend to hold provided population substructure is accounted for using methods, which are now standard^4–6^. Researchers may also run genetic association studies in stratified samples ^1,2^ to reduce the non-genetic variation in the trait and improve statistical power to detect variants not acting through a disease mechanism. However, performing an association test between a genetic variant and a continuous trait in a sample that is stratified, depleted or enriched for a disease outcome (collider) that is associated with both the genetic variant and the trait can result in paradoxical observations (**Figure 1**). Such bias is termed index event bias (IEB) or collider-stratification bias^7^.

**Figure 1.**
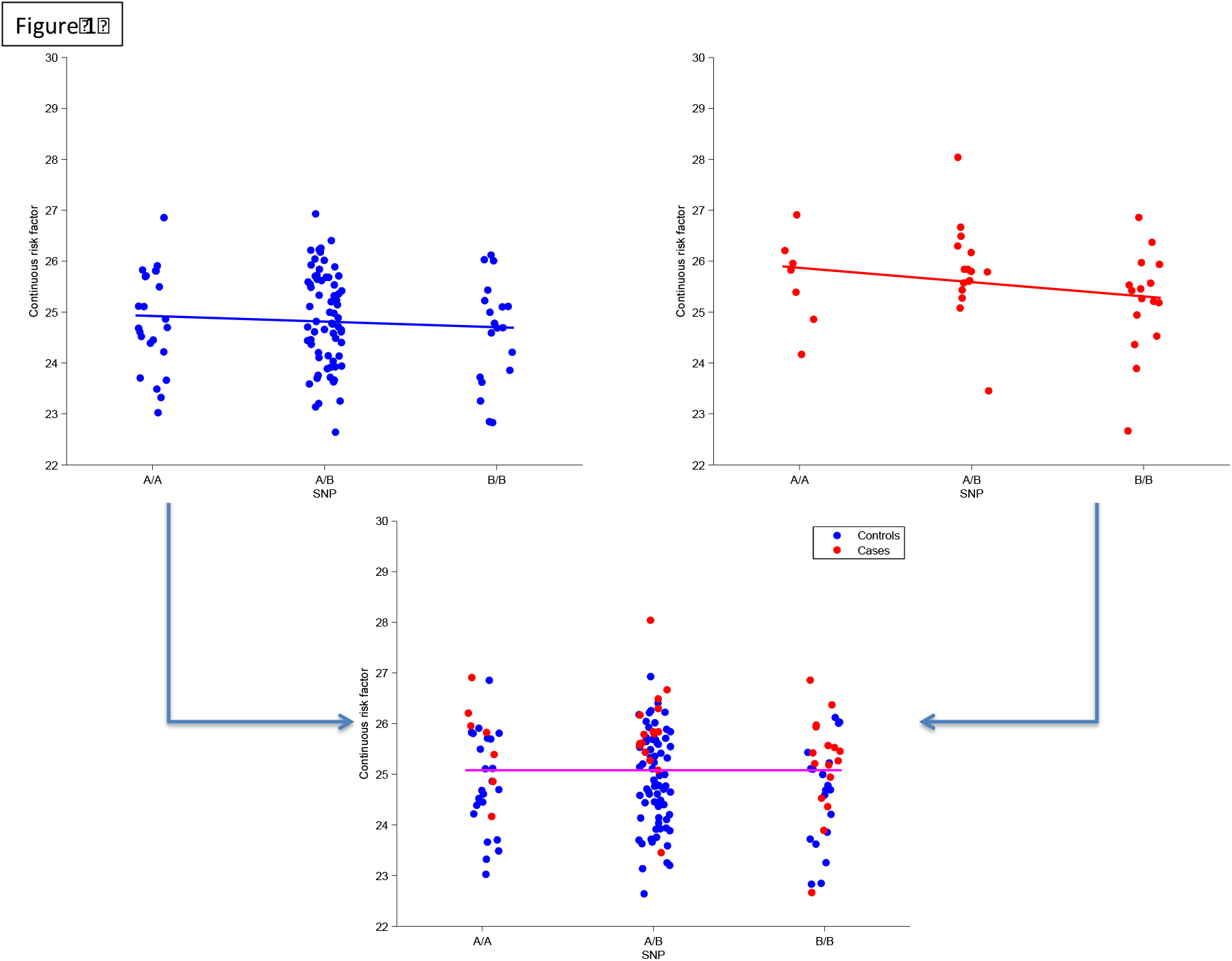
Apparently paradoxical gene-phenotype associations in the context of disease stratified genetic studies. We simulated genotype, continuous risk factor values and disease status in a general population sample according to our liability scale model and set the genetic effect on the risk factor (*γ*) to zero. We observed that the estimated effect of the “B” allele of a genetic marker on a continuous trait is negative both in cases and controls. Disease carriers also have higher trait value than controls. However, when combining the two strata the marker is – as expected – not associated with the trait. The reason for this apparent paradox is that the proportion of disease risk allele (“B”) carriers is higher in the case group. Thus when merging cases into the control group the mean trait value of the BB group increases much more than it does in the other genotype groups. This concept is recognized as Simpson's paradox^40^.

An example of an index event biased association is that between type 2 diabetes risk alleles and lower BMI observed within diabetes case or control strata^8–10^. In these examples the strata are enriched (type 2 diabetes cases) or depleted (type 2 diabetes controls) by disease status. Bias occurs because non-diabetic individuals with a diabetes protective allele are able to remain normo-glycaemic at higher BMIs than individuals without the protective allele, whilst individuals with a risk allele will tend to develop diabetes at lower BMIs compared to those without the risk allele. Considering this type of bias is very important because many large meta-analytic studies often perform GWAS analyses of traits in samples stratified by, enriched or depleted for disease status. Such bias can have different impacts on genetic associations and their interpretation, for example (a) in the case of true (positive) pleiotropy, the effect of the gene variant on the trait may be masked/reduced by IEB (false negative finding); (b) in the case of no pleiotropy, a false (and often counterintuitive) association between gene variant and trait can be observed (false positive finding); (c) using a genetic variant as an instrumental variable in a Mendelian randomization analysis could result in false inferences about causal relationships between the disease and a correlated trait if estimates of that variants’ effects on the trait are biased.

A research area closely related to IEB is secondary trait analysis in case-control sample design. Several approaches have been proposed in these settings to estimate the true effect between a risk factor and a genetic variant in the general population sample^11–16^. Most of these methods yield unbiased estimators that are robust to model misspecifications and work for a wide range of noise distributions. In this paper we applied the SPREG program^16^ to correct for IEB for real data. Note, however, these methods provide only effect size estimation, but no insight how the bias explicitly depends on parameters such as (a) the extent of case-enrichment/depletion; (b) the strength of associations between the SNP, the risk factor and the collider; and (c) SNPs’ allele frequency.

In this study, we tested the extent to which large genetic association studies may be impacted by IEB due to inadvertent sample selection leading to enrichment or depletion of disease cases. We first provided the statistical framework for quantifying IEB in a study, then used a combination of real and simulated data to: (a) identify and quantify real examples of IEB, including a single large study (120,000 individuals from the UK Biobank) and a meta-analysis of independent studies and (b) demonstrate that IEBs can occur in most types of genetic association study designs, e.g. 1:1 case-control designs to case only and control only studies, case or control enriched studies and when case status is used as a covariate. The analytical formula is implemented in an R package and available for download from http://wp.unil.ch/sgg/files/2016/01/IndexEventBias.zip.

## Material and Methods

### Model formulation

We assumed a liability scale disease model, where a genetic variant (*G*) and a continuous risk factor (*X*) influences the development of a disease (*Y*). For simplicity the genetic marker was modeled by a binomially distributed random variable *G* ~ *B*(2,*q*), where *q* is the allele frequency of the genetic variant in the general population. The second trait, from now on referred to as the “continuous risk factor” (*X*) was allowed to be associated with *G* and assumed to follow a standard normal distribution (*X* | *G* = *g*)~ *N*(*γ*(*g* – 2*q*), 1 – 2*γ*^2^*q*(1 – *q*)). The disease status is then modeled as

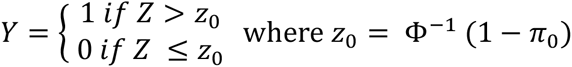

with

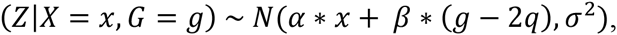

where *π*_0_ is the disease prevalence in the general population; *α* is the true effect size on the liability scale of the continuous risk factor (*X*) on the disease outcome (*Y*); *β* is the (risk-factor independent) effect of the genetic variant (*G*) on the disease outcome (*Y*); and *σ*^2^ = 1 – *α*^2^ – 2*β*^2^*q*(1 – *q*).

### Link between liability scale and logistic models

For simplicity, we derived the explicit analytical formula for IEB estimation for the liability scale model. This however does not prevent its applicability to parameters derived from the logistic regression model. By re-parameterizing the models one can reach indistinguishable properties^17^. Namely, we calculate the probability of (*Y*=0 | *X, G*) for data simulated from the logistic model and optimize the liability scale model parameters such that this probability is matched as close as possible for each simulated data point. We noticed that the models are indistinguishable when the parameter optimization is done for a simulated population with similar disease prevalence to the tested sample. The details of this procedure are given in the Appendix (section 2.3).

### Analytical formula for IEB

The extent of IEB can be analytically derived in the case of the liability scale (probit) disease model. Assume that disease frequency in the general population is *π*_0_, the allele frequencies of the genotype in pure control and pure case populations are *q*_0_ and *q*_1_, respectively. We denote the difference between these frequencies by *δ*_*q*_. Let us consider now a sample with *π* frequency of cases. The allele frequency in this sample is

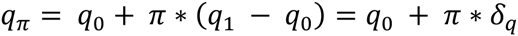

IEB occurs when this disease prevalence differs from the general population prevalence (*π*_0_). When *π* > *π*_0_, we observe an enrichment of cases, while in case of *π* < *π*_0_, we observe an enrichment of controls. In the Supplementary Text we derived the per-allele linear regression effect size of *G* on *X* in the general case, but here for simplicity we present the formula assuming *γ* = 0, i.e. that the true underlying effect of the genotype on the risk factor is zero. We observed that this simplification makes very little difference in practice (**Supplementary Figure 1**). By introducing the quantities

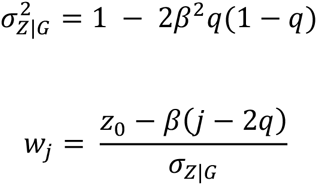

we can express the expectation of the linear regression effect size estimate of *G* on *X* as

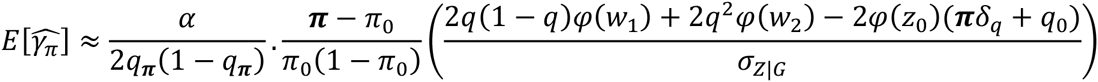

where *φ*() denotes the probability density function (pdf) of the standard Gaussian distribution. Note that when the disease prevalence matches the general population prevalence (i.e. *π* = *π*_0_), the expected effect of *G* on *X* is unbiased. The formula also shows that the bias is a second order rational function of the prevalence (π). For most settings the quadratic and linear terms of the denominator are small compared to that of the numerator, thus the bias as a function of the fraction of cases (*π*) closely resembles a simple parabolic function. It is worth noting that the coefficient of the quadratic term (in *π*) in the numerator 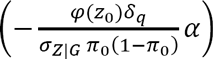 has the opposite sign compared to *α*, meaning that when *X* is a risk factor for the disease the function is a downward looking parabola. This explains why disease risk alleles can show spurious (and counterintuitive) protective effect on traits positively correlated with the disease. We also derived a formula for the case when we do not assume that the true effect is zero; i.e. when there is a true pleiotropy. The full derivation of the formula can be found in the Appendix (Section 2).

Note that this formula assumes that the true parameters, *α*, *β*, *q*_0_, *q*_1_, *q*, *π*_0_ are known. Hence, its primary purpose is not to estimate the bias from data, but to reveal the intricate relationship between the true underlying model parameters and the resulting IEB.

We extended the formula to situations when not only the sample is enriched or depleted for a disease, but also when in addition the continuous risk factor is corrected for the disease status:

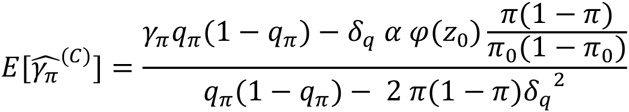

The full derivation of this formula is given in the Appendix (Section 3). One can observe that correcting for disease status yields biased estimates even when the study disease prevalence agrees with the general population prevalence. In this situation we showed that this formula simplifies to that of Aschard^18^.

These analytical formulae for IEB allow us to quantify genetic effects (estimated from real data) for IEB. The key quantities necessary for the formulae are: (i) the allele frequency of the genetic variant in control (*q*_*0*_) and disease (*q*_*1*_) populations; (ii) the association effect of the SNP on the disease status (*β*); (iii) the effect of the continuous risk factor on the disease status (*α*); (iv) the disease prevalence in the study population (*π*) and (v) the general population disease prevalence (*π*_*0*_). Estimating these quantities is out of the scope of this paper. We use the formula to show the extent of the bias for various realistic parameter settings informed from large GWAS data..

### Data simulation to confirm the analytical formula

To investigate how closely the analytical formula recapitulates true IEB, we simulated data to create different hypothetical scenarios similar to data used in genetic studies (**Supplementary Figure 2**). For one million individuals, we simulated binomially distributed SNP data (*G*), a normally distributed continuous risk factor (*X*) and binary disease status using the liability threshold model described above. The minor allele frequency (MAF) of the genetic marker was explored in the range of 0.02, 0.05, 0.1, 0.3 and 0.5; disease prevalence in the general population was set to 1%, 5% and 10%. The effect of the continuous risk factor (*X*) on the disease outcome (*Y*) was varied in a range equivalent to ORs of 1.10 to 4 (1.10, 1.25, 1.50, 1.75, 2, 3 and 4) per 1 standard deviation (SD) increase in *X*. The effect (*β*) of the genetic variant (*G*) on the disease outcome (*Y*) was explored for a range equivalent to odds ratios (OR) of 1.05, 1.2, 1.3, 1.5, 1.8 and 2.

Within each simulated population, we randomly sampled 100,000 controls (individuals with *Y* = 0) and ran a linear regression test between the genetic variant *G* and the continuous risk factor *X*. We then performed 100 iterations each time replacing 1,000 random controls (individuals with *Y* = 0) with 1,000 random cases (individuals with *Y* = 1) and ran a linear regression test between *G* and *X*. Using this approach, our first iteration represented a control-only population of 100,000 unaffected individuals, the 50^th^ iteration a case-control population of 50,000 controls vs. 50,000 cases and the final, 100^th^ iteration a case-only population of 100,000 affected individuals.

### IEB in real data

To identify and quantify real examples of IEB, we tested how IEB occurs in genetic studies in 2 different scenarios using different disease outcomes and genetic variants.

1. We tested whether or not type 2 diabetes risk alleles, acting predominantly through insulin secretion, have a paradoxical (opposite association) effect on BMI, a strong continuous risk factor for type 2 diabetes. We selected 11 SNPs associated with type 2 diabetes that have known robust associations with insulin secretion, the intermediate trait most relevant to diabetes risk ^19^ (**Supplementary Table 1**). We analysed the 11 SNPs separately and as a genetic risk score (GRS) in two study types: (i) a single very large population based study: 120,286 individuals from the first release of genetic data from the UK Biobank study^20^ and (ii) a meta-analysis of 4 independent studies: EXTEND^21^ (N = 5,097), GoDARTS^22^ (N = 7,128), Generation Scotland Scottish Family Health Study (GS:SFHS)^23^ (N = 8,195) and ARIC^24^ (N = 9,324) with a range of study designs and diabetes status available (**Supplementary Table 2**). We tested the association between individual SNPs and the 11-SNP insulin secretion GRS and BMI in all samples, in all samples adjusted for type 2 diabetes status, in diabetic cases only and in controls only. We defined type 2 diabetes cases as individuals who had: (1) HbA1c >6.4% and/or fasting glucose >7 mmol/L, (2) age at diagnosis >35 and <70 years; (3) no need for insulin treatment within 1 year of diagnosis (except in ARIC). We defined controls as individuals who did not meet any of these criteria and were not diagnosed with any other types of diabetes. We additionally tested the association between the *CCND2* type 2 diabetes protective allele and BMI as the allele was recently shown to be associated with higher BMI.
2. We tested whether or not 26 SNPs robustly associated with systolic blood pressure have a paradoxical (opposite) effect on BMI, a continuous risk factor for high blood pressure^25^. We excluded the variant near *SLC39A8* from the GRS as this variant is directly associated with several traits including BMI^26^ and HDL-cholesterol^27^ levels (**Supplementary Table 3**). We analysed the 25 SNPs individually and as a GRS in two study types: (i) a single very large study: 120,286 individuals from the first release of genetic data from the UK Biobank study^20^ and (ii) a meta-analysis of 4 independent studies with blood pressure available: GoDARTS^22^ (N = 6643), Generation Scotland Scottish Family Health Study (GS:SFHS)^23^ (N = 8195), ARIC^24^ (N= 9,290) and BRIGHT^28^ (N = 1808) (**Supplementary Table 2**). We tested the association between the individual SNPs and the 25-SNP blood pressure GRS and BMI in all samples, in all samples adjusted for hypertension status, in hypertensive cases only and in normotensive controls only. We defined hypertensive cases as individuals with systolic blood pressure ≥ 140 mmHg or diastolic blood pressure ≥ 90 mmHg or report use of antihypertensive medications. We defined normotensive controls as individuals with systolic and diastolic blood pressure below these thresholds, and not on medications.

### Statistical analysis in real data

In the relevant studies, we corrected BMI for age and sex and other covariates (for the UK Biobank study this included five within UK genetic principal components, genotyping platform, study center in the UK Biobank study). Residuals were inverse-normal transformed. In each study, we generated genetic risk scores (GRS) by calculating the number of disease risk alleles carried by each individual. We then combined the association results using fixed-effects inverse variance-weighted meta-analysis. To account for IEB we applied the state-of-the-art method of Lin and Zeng^16^ (implemented in the software SPREG). This program needs an estimate of the general population prevalence of the examined diseases. Hence, we derived an estimate for type 2 diabetes and hypertension prevalence for a general UK subpopulation that has the same joint age- and sex-distribution as the UK Biobank sample. For this we used sex- and age-group-specific prevalence values from the IDF Atlas ^29^ (10 year bins) for type 2 diabetes and from the NIH Health Survey for England 2011 [http://digital.nhs.uk/catalogue/PUB09300/HSE2011-Ch3-Hypertension.pdf] (10 year bins) for hypertension. Then we weighted these prevalence values with the proportion of UK Biobank participants that fell into each stratum. This yielded prevalence estimates of 10.15% for type 2 diabetes and 38.43% for hypertension.

## Results

### IEB occurs in a range of different genetic association study designs – theory and simulated data

Our results from simulated data and theory provided examples of IEBs where the direction of the association between the genetic risk allele and the risk factor depends on the proportion of cases and controls in the study. In all scenarios involving a disease risk allele with no real association to the intermediate risk factor, we observed U-shaped artefactual effect estimates between the disease risk allele and the risk factor as the proportion of cases moved from 0% to 100%: first, in a control only situation, IEB occurred where the disease risk alleles were associated with lower values of the risk factor, there was then no association when the proportion of cases represented exactly the background population, then an association between disease risk alleles and higher values of the risk factor, and then back to no association and finally an association between disease risk alleles and lower values of the risk factor in case only scenarios (**Figure 2** [our theoretical formula]; **Supplementary Figure 3** [simulated data]). The extent of the bias is stronger in case only compared to control only scenarios when the disease frequency is less than 50% (as with most diseases). In the examples in **Figure 2** (and **Supplementary Figure 3**), we modeled a disease risk allele and a protective allele with properties similar to those of the type 2 diabetes alleles at *TCF7L2* and *CCND2* respectively. We observed spurious associations between the disease alleles and lower and higher values of the continuous risk factor, depending on the proportion of cases and despite the lack of a genuine association between the genetic risk allele and the risk factor. When the examined study population matches the underlying general population in terms of disease prevalence (5% in case of our example), no bias is observed (**Figure 2**). It has been shown that for many scenarios correcting for disease status alleviates the bias^15^, but such correction is clearly of no use in case-only, and control-only designs and it introduces bias even for samples representative of the general population. Note that in our settings the bias was not resolved, but often exacerbated by correcting for disease status (**Figure 2**).

**Figure 2.**
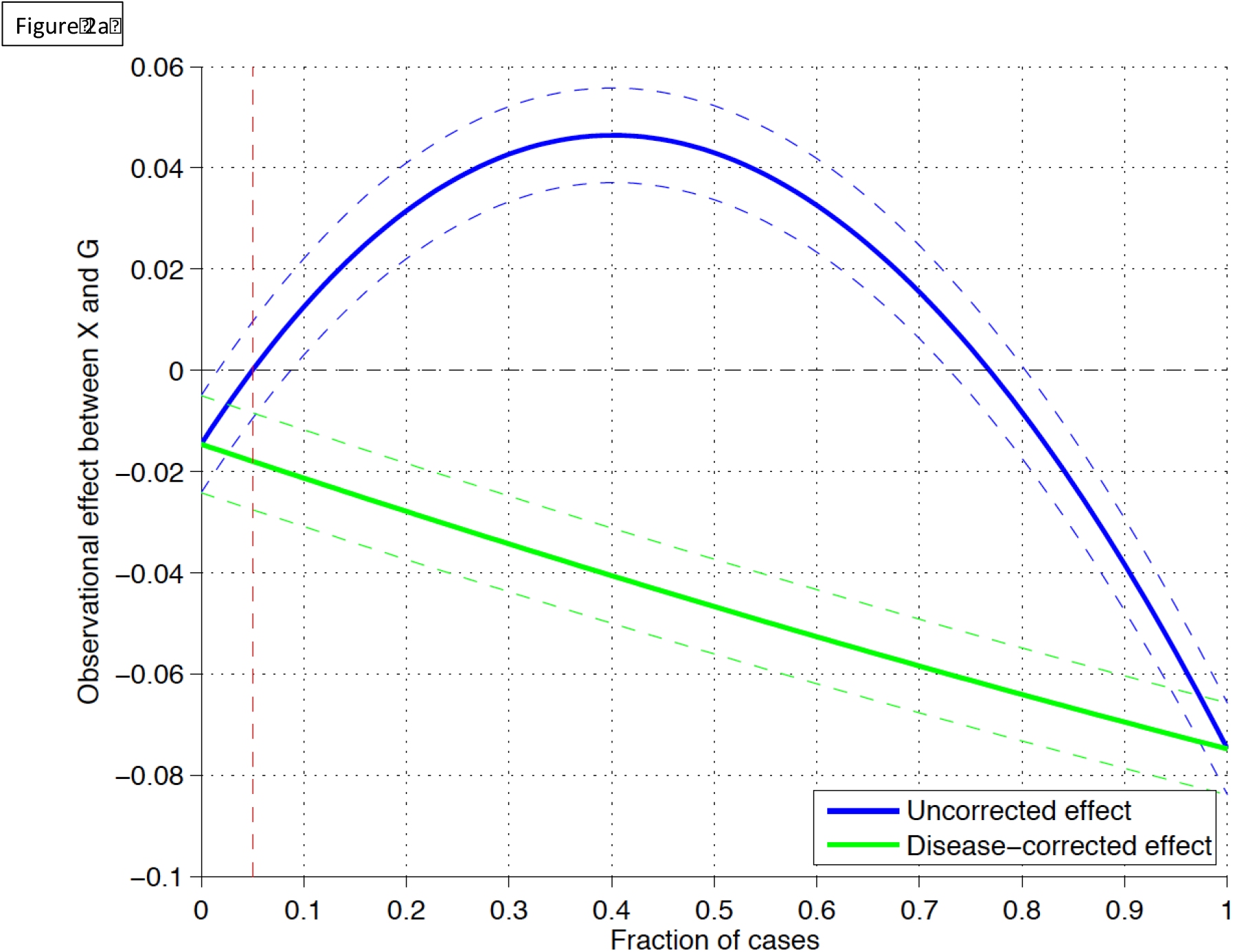
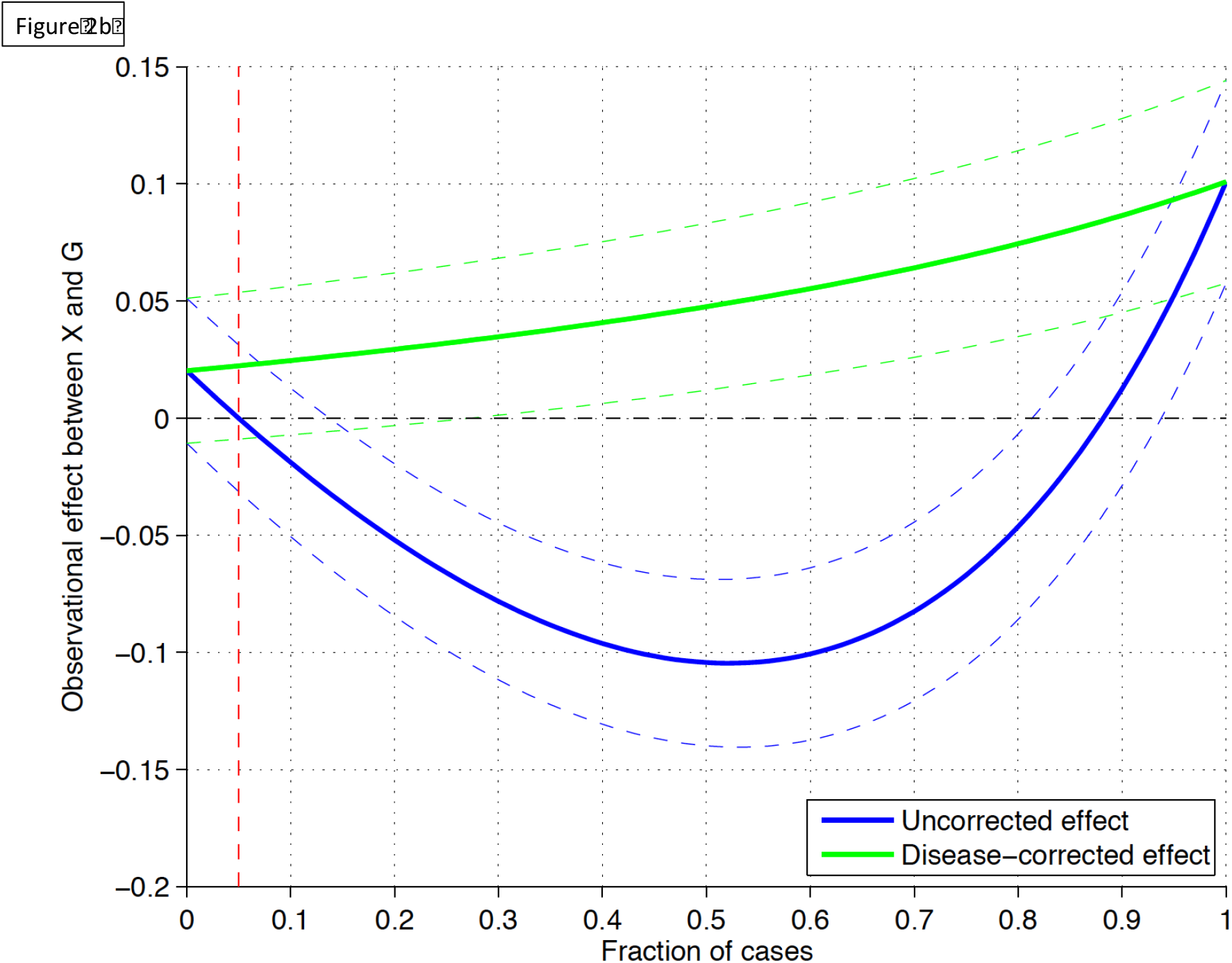
Enrichment for cases or controls produces spurious associations. We applied our analytical formula to compute the effect size estimate of a SNP (*G*) on a continuous risk factor (*X*) in the abovementioned liability scale model setting with the true genetic effect on the risk factor (*γ*) being zero. Enrichment for cases or controls produces spurious evidence of association between disease risk alleles and a risk factor correlated with the disease (equivalent to 2.5 OR per SD) in (**a**) a scenario where a risk allele (MAF 30%) increases risk with an effect equivalent to an odds ratio of 1.4 (similar to the *TCF7L2* type 2 diabetes scenario^10^) in two models: unadjusted for disease status [blue curve] and adjusted for disease status [green curve]. Dashed lines represent 95% confidence interval (CI) around the effect estimate assuming a population of 100,000 individuals. Panel (**b**) displays the same curves, but for a SNP with a rare protective allele (MAF 2%) that reduces risk of disease with an effect equivalent to an odds ratio of 0.5 (similar to the *CCND2* type 2 diabetes scenario^9^). Vertical dashed red line at 0.05 indicates the true general population disease prevalence.

### Differences in prevalence for Type 2 diabetes and hypertension in the UK Biobank and general UK population

Index event bias arises due to differences in the (collider) disease prevalence in the study population and the matched general population. Hence, it is crucial to derived accurate estimates for type 2 diabetes and hypertension prevalence for a general UK sub-population that has the same joint age- and sex-distribution as the UK Biobank. Using data from the IDF Atlas ^29^ and the NIH Health Survey for England, we estimated that 10.15% and 38.43% of a sex- and age-matched sub-population of the UK would be diabetic and hypertensive, respectively (see Methods). These values clearly differ from the prevalence of type 2 diabetes (3.4%) and hypertension (55.2%) observed in the UK Biobank.

### Individual alleles and genetic risk scores associated with higher risk of type 2 diabetes were associated with lower BMI in real data

A relatively high ability to secrete insulin may lead to a relative protection from type 2 diabetes but may also lead to higher BMI because insulin has anabolic properties. Studies may therefore wish to use common variants associated with insulin secretion to test the role of insulin secretion on BMI. However, there may be a complex relationship because higher BMI increases diabetes risk. IEB will add to the complexity of interpreting potential overlap of genetic associations for these phenotypes. We tested 11 variants associated with type 2 diabetes through an insulin secretion mechanism, for potentially spurious associations with lower levels of the continuous risk factor for type 2 diabetes, BMI. Details of how these variants were associated with type 2 diabetes in UK Biobank and 4 additional studies are given in **Supplementary Table 4**. Using a total of 4,003 type 2 diabetes cases and 113,203 controls from the UK Biobank, two of the 11 variants were associated with lower BMI in all individuals (unstratified and unadjusted), three in controls only, three in cases only and five in all individuals when adjusted for type 2 diabetes status at p<0.05 (**Table 1**). When meta-analysing the UK Biobank and four additional studies in the same analysis design as a GWAS meta-analysis (all individuals together for population based studies, stratified by case control status for case control studies) two type 2 diabetes risk alleles were associated at p<0.05 with lower BMI (**Supplementary Table 5**). In the UK Biobank study, the effect sizes of type 2 diabetes risk alleles with lower BMI were consistent with IEB (**Supplementary Figure 4**). In accordance with our formula, BMI “effect” size estimates were correlated with the effect estimates for type 2 diabetes; r = −0.85 (p=8E-4), −0.87 (p=4E-4) and −0.87 (p=5E-4) in controls, cases and all individuals (adjusted for type 2 diabetes status), respectively (**Supplementary Figure 4**).

**Table 1.** Association between 11 insulin secretion SNPs and BMI (SD) in the UK Biobank study under 5 scenarios. Corrected statistics are those after correcting for index event bias using all 119,688 individuals. Note that the numbers of individuals in the “all” analyses differ slightly because people of uncertain diabetes diagnosis were excluded from the “All adjusted for type 2 diabetes” analysis.

| Variant in or near gene: | All<br>(N=119,688) |  | Controls<br>(N=113,203) |  | Cases<br>(N=4,003) |  | All adjusted for type<br>2 diabetes<br>(N=117,206) |  | All - Corrected<br>statistics<br>(N=119,688) |  |
| --- | --- | --- | --- | --- | --- | --- | --- | --- | --- | --- |
|  | BETA | P | BETA | P | BETA | P | BETA | P | BETA | P |
| *TCF7L2* | -0.024 | 1E-7 | -0.033 | 8E-13 | -0.118 | 1E-7 | -0.036 | 1E-15 | -0.0101 | 0.04 |
| *THADA* | -0.011 | 0.1 | -0.012 | 0.08 | -0.060 | 0.1 | -0.013 | 0.05 | -0.0020 | 0.8 |
| *CDKN2AB* | -0.008 | 0.2 | -0.014 | 0.01 | -0.018 | 0.5 | -0.014 | 0.009 | -0.0013 | 0.8 |
| *SLC30A8* | -0.005 | 0.3 | -0.009 | 0.05 | -0.034 | 0.2 | -0.009 | 0.03 | -0.0008 | 0.9 |
| *CDKAL1* | -0.012 | 0.01 | -0.016 | 5E-4 | -0.048 | 0.03 | -0.018 | 1E-4 | -0.0062 | 0.2 |
| *MTNR1B* | 0.011 | 0.02 | 0.008 | 0.08 | 0.004 | 0.9 | 0.008 | 0.08 | 0.0162 | 0.001 |
| *HHEX* | -0.005 | 0.2 | -0.007 | 0.09 | -0.054 | 0.01 | -0.009 | 0.04 | 0.0005 | 0.9 |
| *GCK* | -0.005 | 0.3 | -0.009 | 0.1 | -0.009 | 0.7 | -0.009 | 0.1 | -0.0005 | 0.9 |
| *PROX1* | -0.001 | 0.8 | -0.002 | 0.6 | -0.002 | 0.9 | -0.002 | 0.6 | 0.0023 | 0.6 |
| *ADCY5* | -0.002 | 0.7 | -0.006 | 0.2 | 0.012 | 0.6 | -0.005 | 0.3 | -0.0014 | 0.8 |
| *DGKB* | -0.003 | 0.5 | -0.005 | 0.2 | -0.008 | 0.7 | -0.005 | 0.2 | 0.0021 | 0.6 |

We next reran the SNP-BMI associations, using the statistical software SPREG, which accounts for IEB^16^. Prior to this correction, the risk allele at *TCF7L2* was associated with lower BMI in all scenarios (the overall population as well as stratified and corrected data -**Table 1**). After correcting for IEB there was no evidence (at p<0.005; p-value corrected for multiple testing) for an association between *TCF7L2* and lower BMI (**Table 1**). In contrast, the type 2 diabetes risk allele at *MTNR1B* was the only allele associated with higher BMI in the overall population (p = 0.02); when accounting for IEB it was even more strongly associated (0.016 SD [0.007, 0.026], P=0.001) (**Table 1, Figure 3**). The type 2 diabetes protective allele in *CCND2* (conferring the strongest effect on type 2 diabetes; 0.59 OR [0.48,0.73]; p = 1E-6) had the strongest effect estimate on BMI (0.06 SD [0.03,0.09]; p = 0.0004), which became much weaker after correcting for IEB (0.003 SD [−0.029,0.035]; p = 0.9).

**Figure 3.**
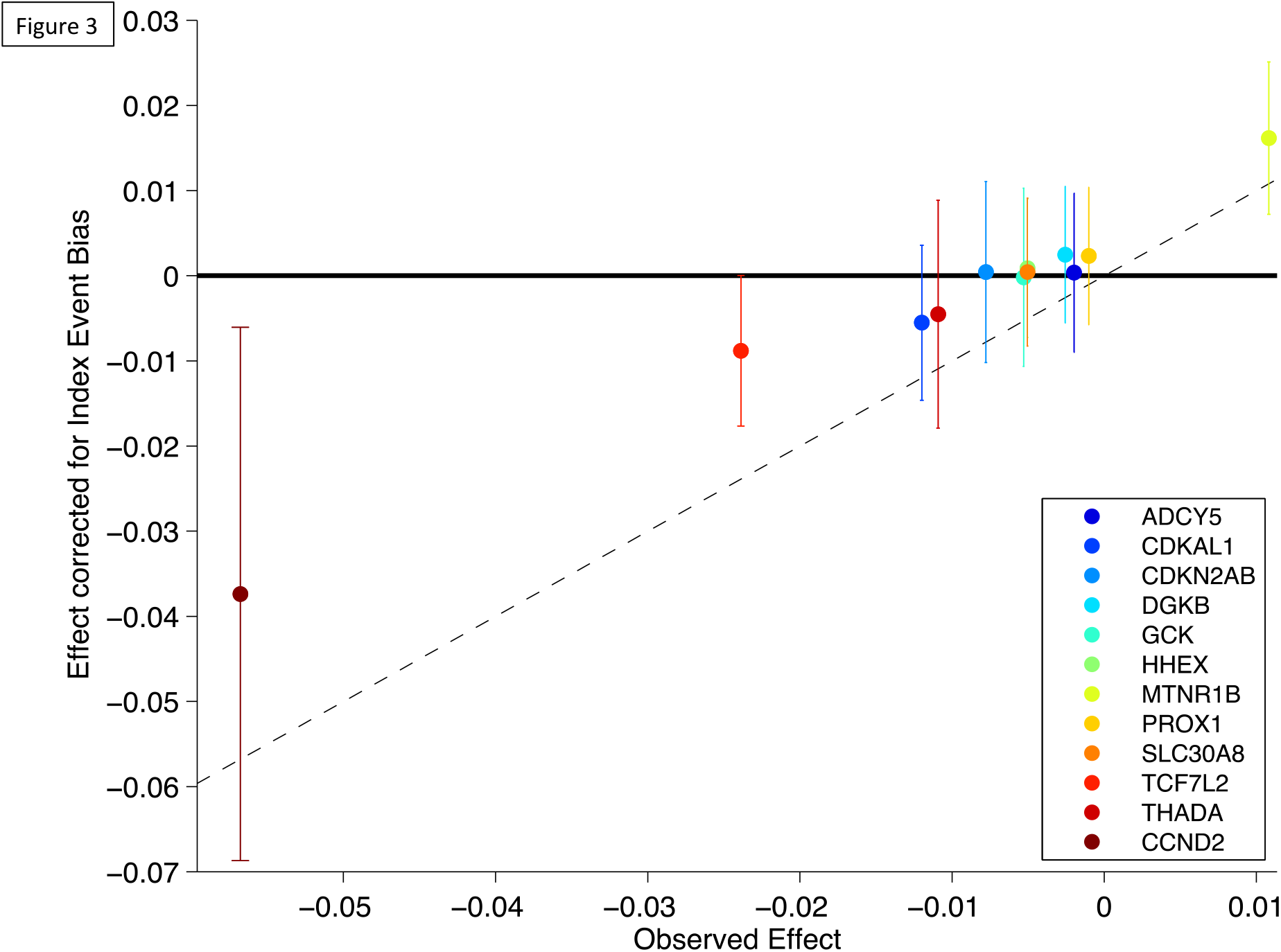
Scatter plot of the observed effect of type 2 diabetes-associated SNPs on BMI in the total UK Biobank sample vs. the index event bias corrected effect. The effect corrected for index event bias (shown on the y-axis) was calculated assuming the previously established 10% population prevalence of type 2 diabetes (*π*_0_ = 0.10). Dashed line represents the identity line, where the two effects are equal. While for most SNPs the absolute value effect size estimate after IEB correction is reduced, *MTNR1B* shows increased effect size upon correction. Only this latter SNP produced a P-value surviving multiple testing correction (P<0.05/11).

We next examined a genetic risk score (GRS) for type 2 diabetes. Details of how this GRS was associated with type 2 diabetes are given in **Supplementary Table 4**. In the UK Biobank study, the 11-SNP GRS was not associated with BMI when analysed in all samples combined (−0.004 SD per allele [−0.010,0.001]; p=0.1; N=119,688; **Table 2**). In contrast, the 11-SNP GRS associated with higher risk of type 2 diabetes was associated with lower BMI in all 3 of the following designs: (i) controls only (−0.010 SD per allele [−0.016,−0.005]; p=5E-4; N=113,203), (ii) in cases only (−0.036 SD per allele [−0.065,−0.007]; p=0.01; N=4,003) and (iii) in all individuals when adjusted for type 2 diabetes status (−0.011 SD per allele [−0.017,−0.006]; p=1E-4; N=117,206; **Table 2**). In the context of a Mendelian randomization analysis, these results could be misinterpreted as evidence for the biologically plausible hypothesis that lower insulin secretion leads to lower BMI. However the associations are consistent with IEB. Results from a meta-analysis of 4 additional studies (representing a scenario similar to that of many GWAS meta-analyses) were similar (**Table 2** and **Figure 4a**).

**Table 2.** Examples of index event bias observed in real data when using multiple variants in genetic risk scores. BMI was inverse-normal transformed.

| Model | Samples | UK Biobank study |  |  |  |  | Meta-analysis of independent studies |  |  |  |  |
| --- | --- | --- | --- | --- | --- | --- | --- | --- | --- | --- | --- |
|  |  | BETA | LCI | UCI | P | N | BETA | LCI | UCI | P | N |
| BMI ~ insulin secretion GRS | All individuals | -0.004 | -0.010 | 0.001 | 0.14 | 119,688 | 0.001 | -0.010 | 0.013 | 0.8 | 30,440 |
|  | Type 2 diabetes controls | -0.010 | -0.016 | -0.005 | 5E-4 | 113,203 | -0.008 | -0.019 | 0.004 | 0.2 | 25,039 |
|  | Type 2 diabetes cases | -0.036 | -0.065 | -0.007 | 0.015 | 4,003 | -0.037 | -0.062 | -0.012 | 0.004 | 5,396 |
|  | All individuals adjusted for type 2 diabetes status | -0.011 | -0.017 | -0.006 | 1E-4 | 117,206 | -0.013 | -0.021 | -0.005 | 0.002 | 30,435 |
| BMI ~ blood pressure GRS | All individuals | -0.014 | -0.020 | -0.008 | 1E-6 | 119,688 | -0.008 | -0.020 | 0.004 | 0.2 | 25,059 |
|  | Normotensive controls | -0.034 | -0.043 | -0.026 | 2E-16 | 53,377 | -0.014 | -0.028 | 0.000 | 0.06 | 18,590 |
|  | Hypertensive cases | -0.031 | -0.039 | -0.024 | 4E-16 | 65,584 | -0.042 | -0.063 | -0.021 | 9E-5 | 8,267 |
|  | All individuals adjusted for hypertension status | -0.033 | -0.038 | -0.027 | 7E-31 | 118,961 | -0.022 | -0.034 | -0.010 | 4E-4 | 25,049 |

**Figure 4.**
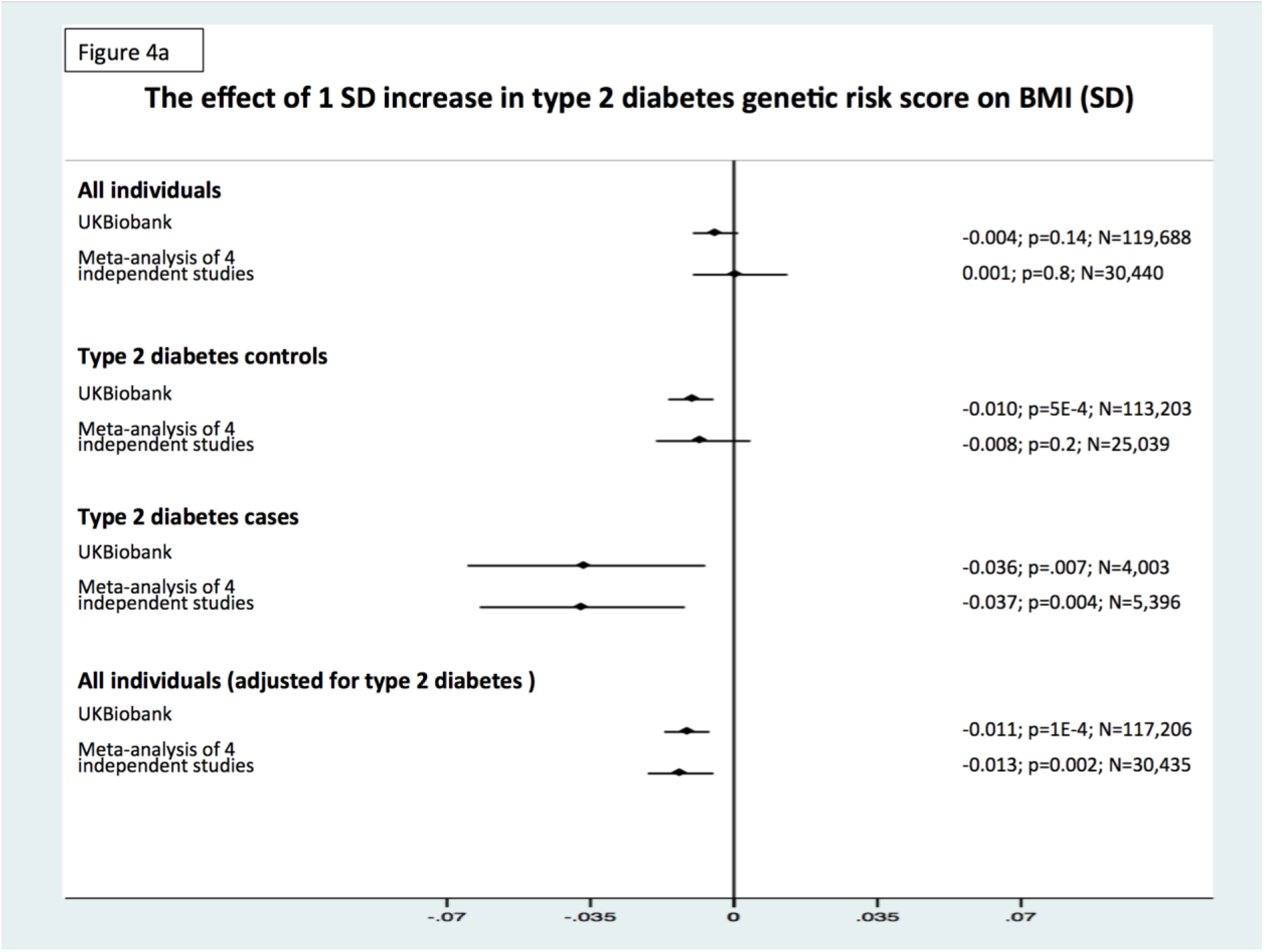
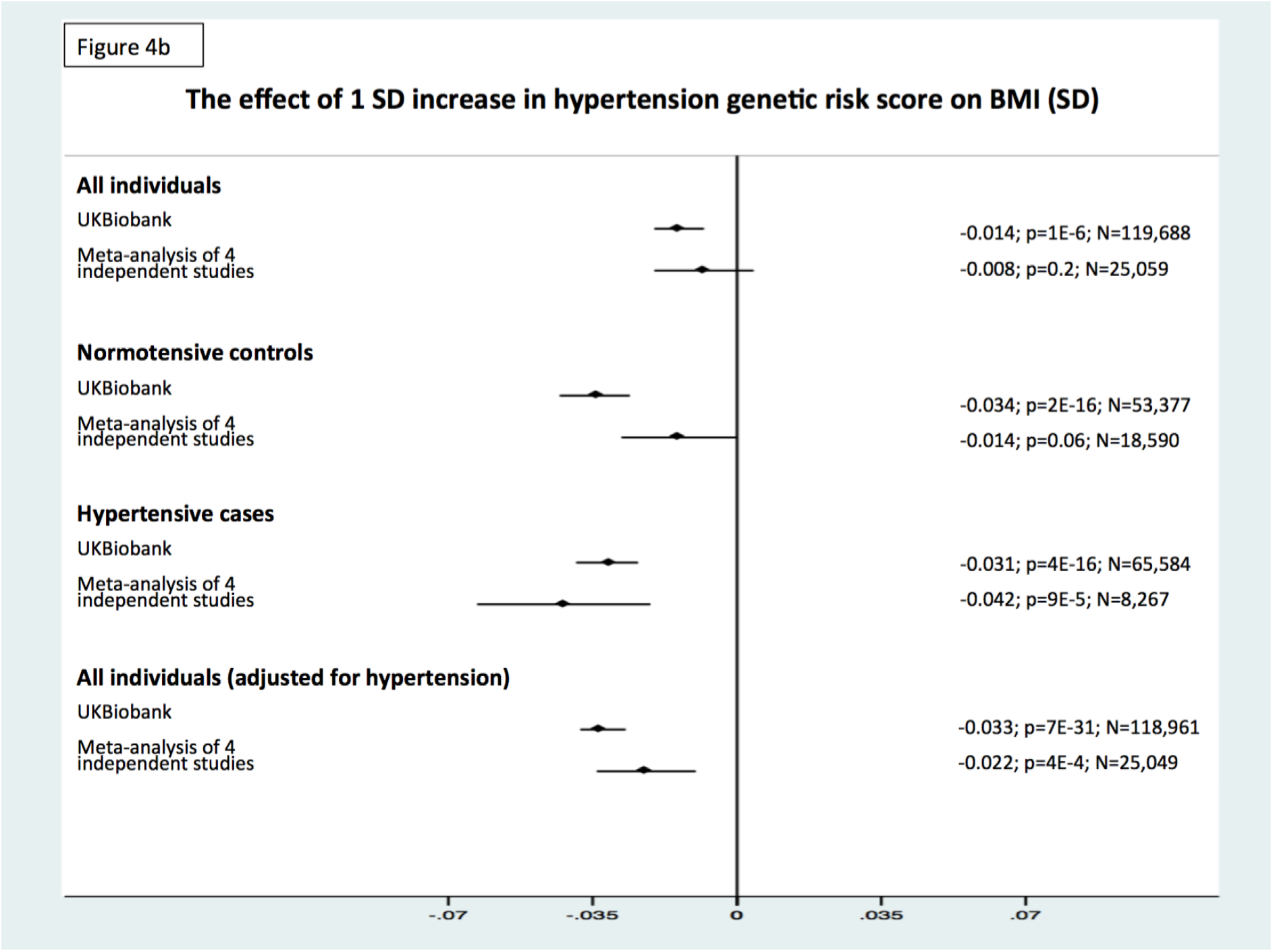
Index event bias in real data. (**a**) The genetic risk score (GRS) associated with higher risk of type 2 diabetes is associated with lower BMI in cases and controls separately and when combined but adjusted for type 2 diabetes status. (**b**) The GRS associated with higher risk of hypertension is associated with lower BMI in hypertensive cases and controls separately and when combined but adjusted for hypertension status. The x-axis is the effect size per disease risk allele. The vertical solid line is the null effect.

### Individual alleles and genetic risk scores associated with higher risk of hypertension were associated with lower BMI

We next tested whether alleles associated with higher risk of hypertension were paradoxically associated with lower BMI, a continuous risk factor for hypertension, but with a weaker effect than that with type 2 diabetes. Such associations could be due to genuine pleiotropic effects of alleles on hypertension and lower BMI, or due to IEB, or a combination of the two. We tested 25 variants associated with blood pressure. Details of how these variants were associated with hypertension in UK Biobank and four additional studies are given in **Supplementary Table 6**. Using a total of 65,584 hypertension cases and 53,377 controls from the UK Biobank, six of the 25 variants were associated with lower BMI in all individuals (unstratified and unadjusted), 10 in controls only, 10 in cases only and 12 in all individuals when adjusted for hypertension status (at p<0.05; **Table 3**). When meta-analysing the UK Biobank and four additional studies, in the same analysis design as a GWAS meta-analysis (all individuals together for population based studies, stratified by case control status for case control studies), eight hypertension risk alleles were associated at p<0.05 with lower BMI (**Supplementary Table 7**). The effect sizes of hypertension risk alleles with lower BMI were consistent with IEB. As with type 2 diabetes alleles, BMI “effect” sizes were correlated with the effect size on hypertension: r = −0.38 (p=0.06), −0.58 (p=0.002) and −0.55 (p=0.005) in controls, cases and all individuals (adjusted for hypertension status), respectively (**Supplementary Figure 5**).

**Table 3.** Association between 25 blood pressure SNPs and BMI (SD) in the UK Biobank study under 5 scenarios. Corrected statistics are those after correcting for index event bias using all 119,688 individuals. Note that the numbers of individuals in the “all” analyses differ slightly because people of uncertain hypertension diagnosis were excluded from the “All adjusted for hypertension” analysis.

|  | All<br>(N=119,688) |  | Controls<br>(N= 53,377) |  | Cases<br>(N=65,584) |  | All adjusted for<br>hypertension<br>(N=118,961) |  | All - Corrected<br>statistics<br>(N=119,688) |  |
| --- | --- | --- | --- | --- | --- | --- | --- | --- | --- | --- |
| Variant in or<br>near gene: | BETA | P | BETA | P | BETA | P | BETA | P | BETA | P |
| *ADM* | 0.001 | 0.9 | 0.005 | 0.6 | -0.012 | 0.1 | -0.005 | 0.4 | 0.0009 | 0.9 |
| *ATP2B1* | -0.002 | 0.7 | -0.012 | 0.1 | -0.011 | 0.1 | -0.012 | 0.03 | -0.0038 | 0.5 |
| *BAT2* | -0.013 | 2E-3 | -0.014 | 0.02 | -0.016 | 3E-3 | -0.015 | 2E-4 | -0.0135 | 0.002 |
| *C10orf107* | -0.008 | 0.1 | -0.012 | 0.1 | -0.019 | 9E-3 | -0.016 | 3E-3 | -0.0095 | 0.08 |
| *CACNB2(3')* | -0.01 | 0.03 | -0.02 | 1E-3 | -0.013 | 0.02 | -0.016 | 1E-4 | -0.0100 | 0.02 |
| *CACNB2(5')* | -0.012 | 4E-3 | -0.023 | 1E-4 | -0.01 | 0.06 | -0.016 | 8E-5 | -0.0122 | 0.003 |
| *CYP17A1* | -0.028 | 3E-4 | -0.044 | 4E-5 | -0.032 | 2E-3 | -0.038 | 5E-7 | -0.0282 | 0.0003 |
| *CYP11A1* | -0.011 | 0.01 | -0.017 | 9E-3 | -0.016 | 7E-3 | -0.016 | 2E-4 | -0.0110 | 0.01 |
| *EBF1* | -0.001 | 0.89 | -0.012 | 0.04 | -0.002 | 0.7 | -0.007 | 0.1 | -0.0008 | 0.9 |
| *FGF5* | -0.007 | 0.1 | -0.023 | 4E-4 | -0.014 | 0.01 | -0.018 | 3E-5 | -0.0078 | 0.08 |
| *FLJ32810* | -0.008 | 0.07 | -0.014 | 0.03 | -0.016 | 8E-3 | -0.015 | 6E-4 | -0.0084 | 0.07 |
| *FURIN* | 0 | 0.98 | -0.008 | 0.2 | -0.008 | 0.2 | -0.008 | 0.07 | -0.0005 | 0.9 |
| *GNAS* | -0.004 | 0.5 | -0.007 | 0.5 | -0.019 | 0.02 | -0.014 | 0.02 | -0.0042 | 0.5 |
| *GOSR2* | -0.001 | 0.8 | -0.011 | 0.2 | -0.003 | 0.7 | -0.007 | 0.2 | -0.0012 | 0.8 |
| *HFE* | 0.002 | 0.8 | -0.008 | 0.3 | -0.001 | 0.9 | -0.004 | 0.5 | 0.0026 | 0.6 |
| *JAG1* | 0 | 0.95 | -0.005 | 0.4 | -0.002 | 0.8 | -0.003 | 0.4 | -0.0005 | 0.9 |
| *MECOM* | 0.002 | 0.7 | -0.006 | 0.3 | 0.001 | 0.9 | -0.002 | 0.6 | 0.0016 | 0.7 |
| *MOV10* | 0.007 | 0.1 | 0 | 0.97 | 0.007 | 0.3 | 0.004 | 0.4 | 0.0079 | 0.1 |
| *MTHFR* | -0.011 | 0.05 | -0.018 | 0.02 | -0.024 | 1E-3 | -0.021 | 7E-5 | -0.0110 | 0.05 |
| *NPR3* | 0.004 | 0.3 | -0.001 | 0.9 | -0.006 | 0.3 | -0.004 | 0.4 | 0.0042 | 0.3 |
| *PLCE1* | -0.001 | 0.8 | 0 | 0.98 | -0.009 | 0.1 | -0.005 | 0.2 | -0.0017 | 0.7 |
| *PLEKHA7* | -0.005 | 0.3 | -0.016 | 0.01 | 0.002 | 0.8 | -0.006 | 0.2 | -0.0049 | 0.3 |
| *SH2B3* | -0.009 | 0.03 | -0.008 | 0.2 | -0.023 | 2E-5 | -0.016 | 4E-5 | -0.0093 | 0.02 |
| *TBX5* | -0.003 | 0.6 | -0.004 | 0.6 | -0.008 | 0.2 | -0.006 | 0.2 | -0.0031 | 0.5 |
| *ZNF652* | 0.006 | 0.1 | 0 | 0.98 | 0.005 | 0.4 | 0.003 | 0.5 | 0.0063 | 0.1 |

Using SPREG, out of the 25 hypertension SNPs only *CYP17A1* was associated with lower BMI in the IEB corrected analysis (−0.028 [−0.044,−0.012]; P=3E-4) and Bonferroni correction for the number of SNPs tested (**Table 3**). Five other variants (those in or near *BAT2, CACNB2* (2 variants)*, CYP1A1,* and *SH2B3*) were associated with lower BMI at p<0.05 but did not persist after Bonferroni correction. Nevertheless, six variants reaching IEB-corrected nominally significant P-values is more than the ~1 expected by chance (enrichment P = 1.69E-4) and suggests variants in some of these genes have pleiotropic effects with alleles associated with lower BMI and higher risk of hypertension. Consistent with this evidence of pleiotropy, the variant in *SH2B3* is associated with multiple traits including those related to autoimmunity as well as metabolic traits^30–32^.

We next considered a genetic risk score of hypertension SNPs. Details of how this GRS was associated with hypertension are given in **Supplementary Table 6**). In the UK Biobank study, the 25-SNP hypertension GRS was associated with lower BMI in all samples combined (−0.014 SD per allele [−0.020,−0.008]; p=1E-6; N=119,688) and in all 3 of the following designs: (i) in controls only (−0.034 SD per allele [−0.043,−0.026]; p=2E-16; N=53,377), in cases only (−0.031 SD per allele [−0.039,−0.024]; p=4E-16; N=65,584) and in all samples when adjusted for hypertension status (−0.033 SD per allele [−0.038,−0.027]; p=7E-31; N=118,961; **Table 2**). Results from a meta-analysis of 4 studies (representing a GWAS meta-analysis) were similar (**Table 2** and **Figure 4b**).

### Large sample sizes are necessary to observe false positive associations due to IEB

Our analytical formula enabled us to quantify the necessary sample size in order to observe a false positive association with a continuous risk factor due to IEB at any significance level alpha with for example, 80% power. The necessary sample size depends on five parameters: significance level, disease prevalence, strength of association between the genetic risk factor and the disease, strength of association between the continuous risk factor and the disease and frequency of the genetic risk allele. We fixed the continuous risk factor-disease association (OR=2.5 per SD) and tested two MAF scenarios (low, 2% and medium 30%) and two significance levels (0.05 and 5E-8). The remaining two parameters (SNP-disease association strength and disease prevalence) we varied freely and computed the minimal sample size necessary to detect a false association (**Figure 5**). For example, in analyses stratified by disease we would need 23,542 cases or 208,267 controls to detect a biased association at p-value 5×10^−8^ with a probability of 80% when the disease risk allele had a frequency of 30%, the disease prevalence was 10% and a 1 SD higher value of the continuous risk factor was associated with an odds ratio of 2.5 for the disease. This scenario is similar to that for the risk allele at *TCF7L2* and BMI^10^.

**Figure 5.**
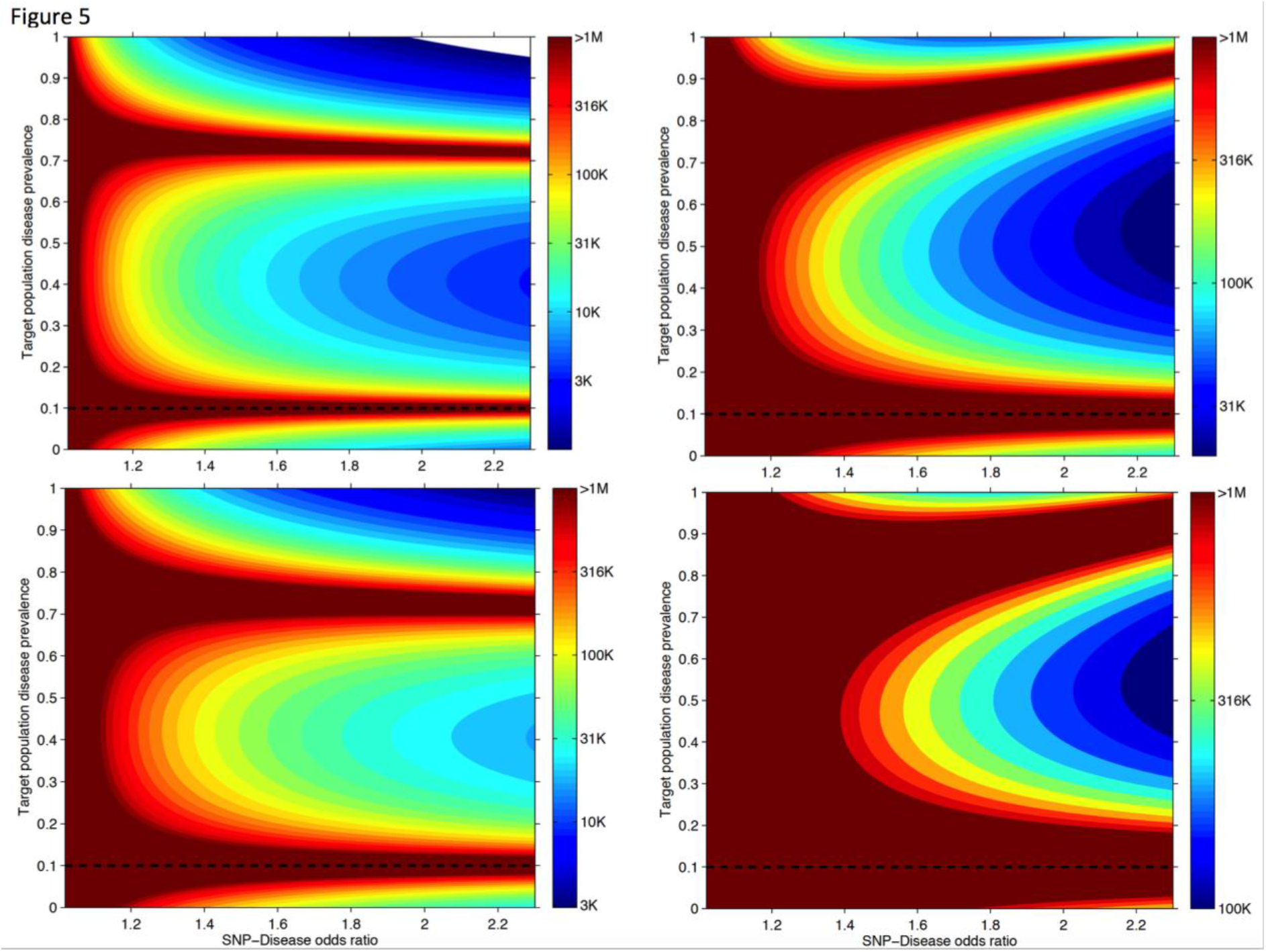
Power calculation to detect IEB. Using the analytical formula for IEB we derived the minimal necessary sample size to observe IEB in a study at nominal (alpha=0.05, top panels) and genome-wide significant level (alpha = 5E-8, bottom panels) with 80% power. We fixed the disease prevalence in the general population to 10%. The SNP-disease odds ratio was varied between 1 and 2.3 and the observed population prevalence of the disease was explored for the full range of 0–100%. The SNP MAF was set to 30% in the left panels and to 2% in right panel.

## Discussion

Our analyses of real and simulated data showed that index event bias will often occur in genetic association studies but the extent depends on several factors. These factors include the association strength between the trait being analysed (that we termed the “continuous risk factor”) and the disease, where the disease is over- or under-represented in the study population compared to the background population. The other factors are disease prevalence, sample size, and the effect size and minor allele frequency of the disease associated variant.

Our results go beyond those of previous studies examining index event biases in several ways. First we provide real examples of likely biased genetic associations in the context of studies of 100,000s of individuals, including those involving individual variants and combinations of variants. Second, we provide a formula for quantifying the bias as a function of key parameters even when only summary level data is provided. We also extended the work of Aschard *et al*^18^, to test how the combination of correcting for disease status in disease-enriched or depleted samples can introduce biases.

Our results have important implications for all types of large genetic association studies, and are especially relevant given that analyses are now possible in 100,000s of individuals, and rarely will these samples be perfectly representative of the background populations – for example, even population based studies such as those of Decode and the UK Biobank are likely not truly representative of the background population in the prevalence of all disease outcomes. Our analyses of real and simulated data showed that the best study design to avoid index event biased associations is using all individuals from a population-based study with no adjustment for disease status. Bias is strongest in case only designs (assuming the disease frequency is <50%) but it is also observed in control only designs, or in analysis combining cases and controls and adjusting for disease status (the latter situation is discussed in Aschard et al^18^). To better understand these observations, we derived an analytical formula that estimates IEB and confirmed its validity through simulation studies. Our formula indicates that this bias can be negative or positive depending on the proportion of cases and controls. The results indicate that this bias cannot be resolved by correcting for disease status as a covariate. If anything, such correction exacerbates biased effect estimates. Our results also indicate that the impact of IEB is substantially larger than the bias caused by improper covariate correction in a disease-representative population (described by Aschard *et al*^18^).

In a meta-analysis it may be difficult to assess the extent to which IEB is contributing to an association - most existing large scale genetic association studies are mixtures of all types of study designs, and, for studies of continuous traits (such as BMI, lipid levels or blood pressure) disease cases and controls are often analysed as separate strata before meta-analysis with population-based studies, which themselves could be over or under represented with disease cases.

Our settings for the analytical formula were limited to a liability scale disease model and normal linear regression applied for the risk trait. By model re-parameterization we extended it to the logistic disease model and through simulations we saw that it works equally well (data not shown). These are the most often used models in meta-analytic GWAS studies; hence we believe that our findings are extremely relevant for almost all GWAS analysis.

Whilst index event biases are likely to exist in many studies, for associations modelling individual variants the bias is unlikely to cause false positive or false negative associations unless sample sizes are very large or stratified, strongly depleted of, or enriched for, disease cases. For example, we tested known common type 2 diabetes variants for association with a strong risk factor for type 2 diabetes, BMI, in 119,688 UK Biobank individuals (including 4003 type 2 diabetes cases), but only the most strongly associated diabetes variant, that in *TCF7L2* (odds ratio ~1.4), was associated with lower BMI at p<0.01 in all individuals. The association between the variant in *TCF7L2* and BMI has been the subject of several previous papers^1,8,33^ and was most recently noted as showing stronger effect estimates in case control studies^1^. Here we found only a trend, but no clear statistical evidence that the *TCF7L2* variant is associated with BMI - the associations we did see could be explained by IEB due to a likely depletion of cases in the UK Biobank study compared to the background population.

The derived analytical formula can serve as guidance for the expected bias in genome-wide association studies, where only summary level data is known where – to our knowledge – no other method is applicable. The state-of-the-art tool, SPREG, computed the corrected effect in ~2 min per SNP, which renders such methods infeasible for large population cohorts with genome-wide genotype data, as it would take >10 CPU years to apply for millions of markers.

Accounting for IEB strengthened associations for several individual variants with good prior evidence for pleiotropic effects on the disease and continuous risk factor. For example, the type 2 diabetes risk allele at *MTN1RB* was associated in the UK Biobank with higher BMI and this result strengthened on correction for IEB – results from previous studies, particularly those that were not population-based, may have been biased towards the null. This variant has one of the strongest effects on fasting glucose levels in individuals without diabetes and may predispose to higher BMI through higher insulin secretion. The hypertension risk allele at *CYP17A1* was previously associated with lower BMI^1^, and we show here that this is a likely pleiotropic effect.

Studies examining the joint effect of multiple variants will be more prone to index event biases than those of single variants. Studies prone to miss-interpretation could include gene-based tests, and tests of the cumulative effect of variants when using a genetic risk score. For example, Mendelian randomization studies often use genetic risk scores as instruments to test causality of an associated trait with a disease^34–37^. Stratified Mendelian randomization is recommended when the exposure is binary (e.g. smoking) and hence a causal effect should be seen only in the exposed stratum^38^. Such stratification in some cases could introduce IEB in the causal estimation^38^. Failing to account for IEB could lead to false conclusions about causality. We explored this potential source of bias using the UK Biobank to assess whether or not a genetic risk score of insulin secretion (represented by 11 variants associated with type 2 diabetes through insulin secretion mechanisms) was associated with BMI. An association between a genetic risk score for poorer insulin secretion and lower BMI could indicate that insulin secretion causally alters BMI, a plausible hypothesis given that insulin treatment increases BMI in diabetes^39^. However, IEB would also result in an association between a genetic risk score for poorer insulin secretion (type 2 diabetes risk alleles) and lower BMI. Whilst we cannot disentangle IEB from a genuine pleiotropic effect IEB is the more likely explanation given the gradient of stronger effects in cases compared to controls compared to all individuals (**Supplementary Figure 6a**). Similar analysis for hypertension provided evidence that SNPs associated with higher blood pressure are also associated with lower BMI (**Supplementary Figure 6b**).

In summary, as genetic association studies reach sizes of 100,000s of individuals, analyses will be prone to misinterpreting results if they do not account for index event biases. However, we have provided the statistical framework and its software implementation for quantifying and correcting for these biases under reasonable assumptions.

## Acknowledgments

This research has been conducted using the UK Biobank Resource. The authors thank University of Exeter Medical School. EXTEND data were provided by the Peninsula Research Bank, part of the NIHR Exeter Clinical Research Facility. P.B.M. wishes to acknowledge support from the NIHR Cardiovascular Biomedical Research Unit at Barts and The London, Queen Mary University of London, UK. We are grateful to all the participants who took part in the GS:SFHS study, to the general practitioners, to the Scottish School of Primary Care for their help in recruiting the participants, and to the whole team, which includes interviewers, computer and laboratory technicians, clerical workers, research scientists, volunteers, managers, receptionists, and nurses. The Wellcome Trust provides support for Wellcome Trust United Kingdom Type 2 Diabetes Case Control Collection (GoDARTS) and informatics support is provided by the Chief Scientist Office. The Atherosclerosis Risk in Communities Study (ARIC) is carried out as a collaborative study supported by National Heart, Lung, and Blood Institute contracts (HHSN268201100005C, HHSN268201100006C, HHSN268201100007C, HHSN268201100008C, HHSN268201100009C, HHSN268201100010C, HHSN268201100011C, and HHSN268201100012C), R01HL087641, R01HL59367 and R01HL086694; National Human Genome Research Institute contract U01HG004402; and National Institutes of Health contract HHSN268200625226C. The authors thank the staff and participants of the ARIC study for their important contributions. Infrastructure was partly supported by Grant Number UL1RR025005, a component of the National Institutes of Health and NIH Roadmap for Medical Research.

H.Y., A.R.W. and T.M.F. are supported by the European Research Council grant: 323195; SZ-245 50371-GLUCOSEGENES-FP7-IDEAS-ERC. S.E.J. is funded by the Medical Research Council (grant: MR/M005070/1). M.A.T., M.N.W. and A.M. are supported by the Wellcome Trust Institutional Strategic Support Award (WT097835MF). R.M.F. is a Sir Henry Dale Fellow (Wellcome Trust and Royal Society grant: 104150/Z/14/Z). R.B. is funded by the Wellcome Trust and Royal Society grant: 104150/Z/14/Z. J.T. is funded by a Diabetes Research and Wellness Foundation Fellowship. Z.K. received financial support from the Leenaards Foundation, the Swiss Institute of Bioinformatics and the Swiss National Science Foundation (31003A-143914) and SystemsX.ch (51RTP0_151019). The work of M.P.B was supported by the National Heart, Lung, And Blood Institute of the National Institutes of Health under Award Number T32HL007779. Generation Scotland received core support from the Chief Scientist Office of the Scottish Government Health Directorates [CZD/16/6] and the Scottish Funding Council [HR03006]. E.R.P. holds a WT New investigator award 102820/Z/13/Z.

We would like to thank George Davey Smith, Mark McCarthy and Joel Hirschhorn for helpful comments on the manuscript.

## Supplementary Figures

**Supplementary Figure 1.**
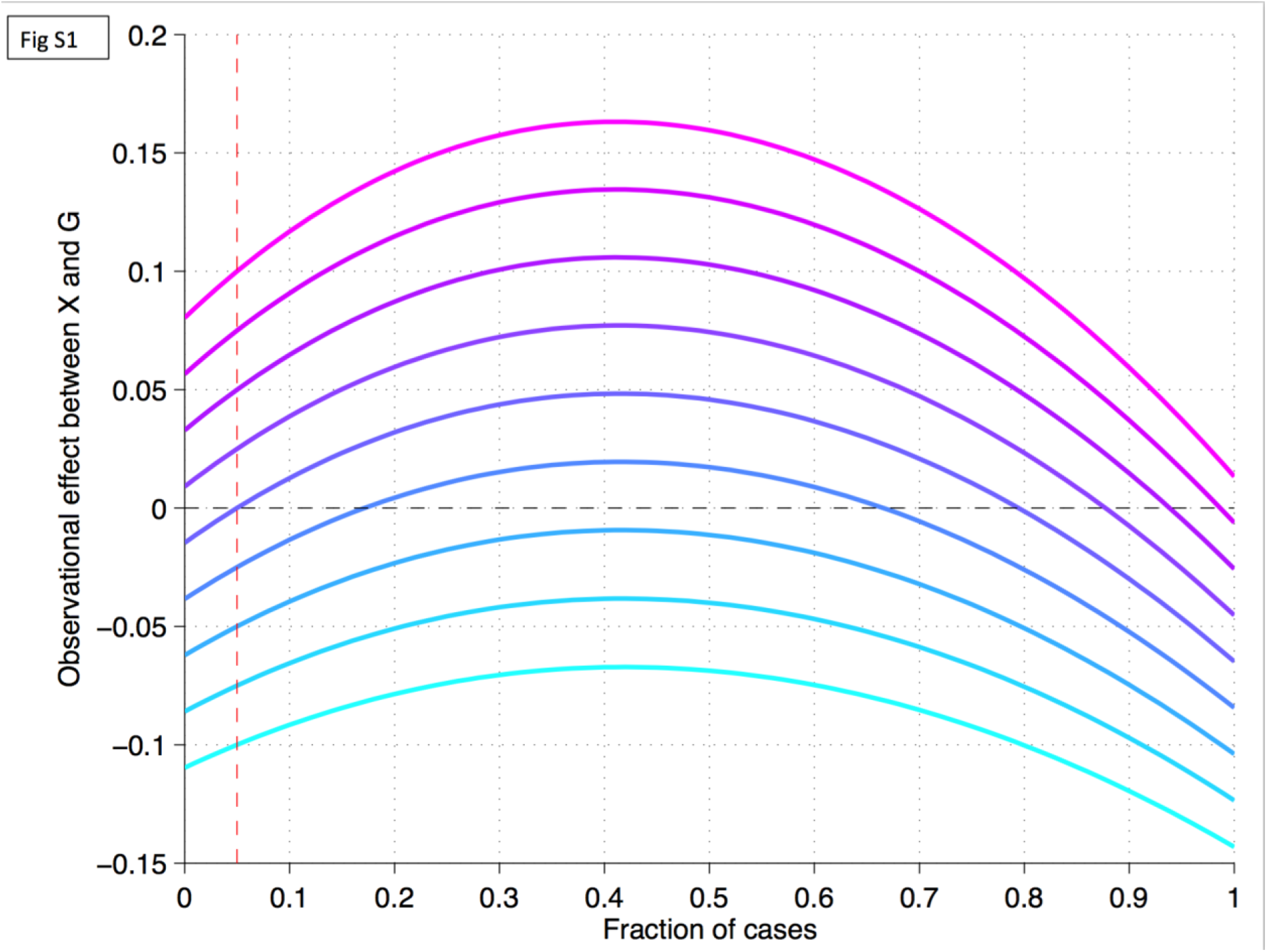
Expected linear regression effects of G on the continuous risk factor using the full index event bias formula. (see Supplementary Text). All notations and settings are identical to the ones in Figure 2, except that here the true (G-X) effect varies between −0.1 and 0.1. One can observe that, in the case-only scenario, the bias is −0.0862, −0.0645 and −0.0429 when the true (G-X) effect is 0.1, 0 and −0.1, respectively.

**Supplementary Figure 2.**
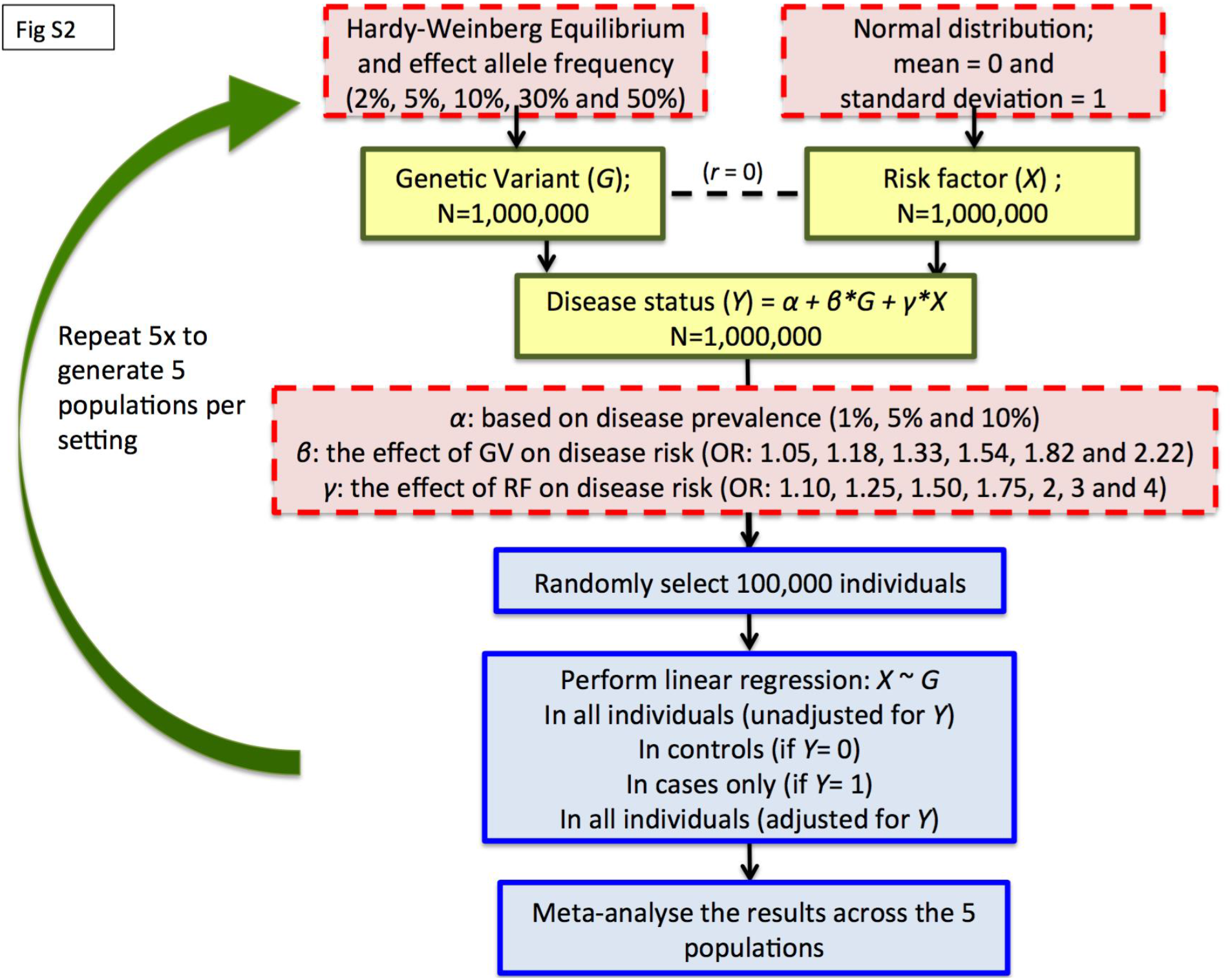
Diagram of study design for the simulation analysis.

**Supplementary Figure 3.**
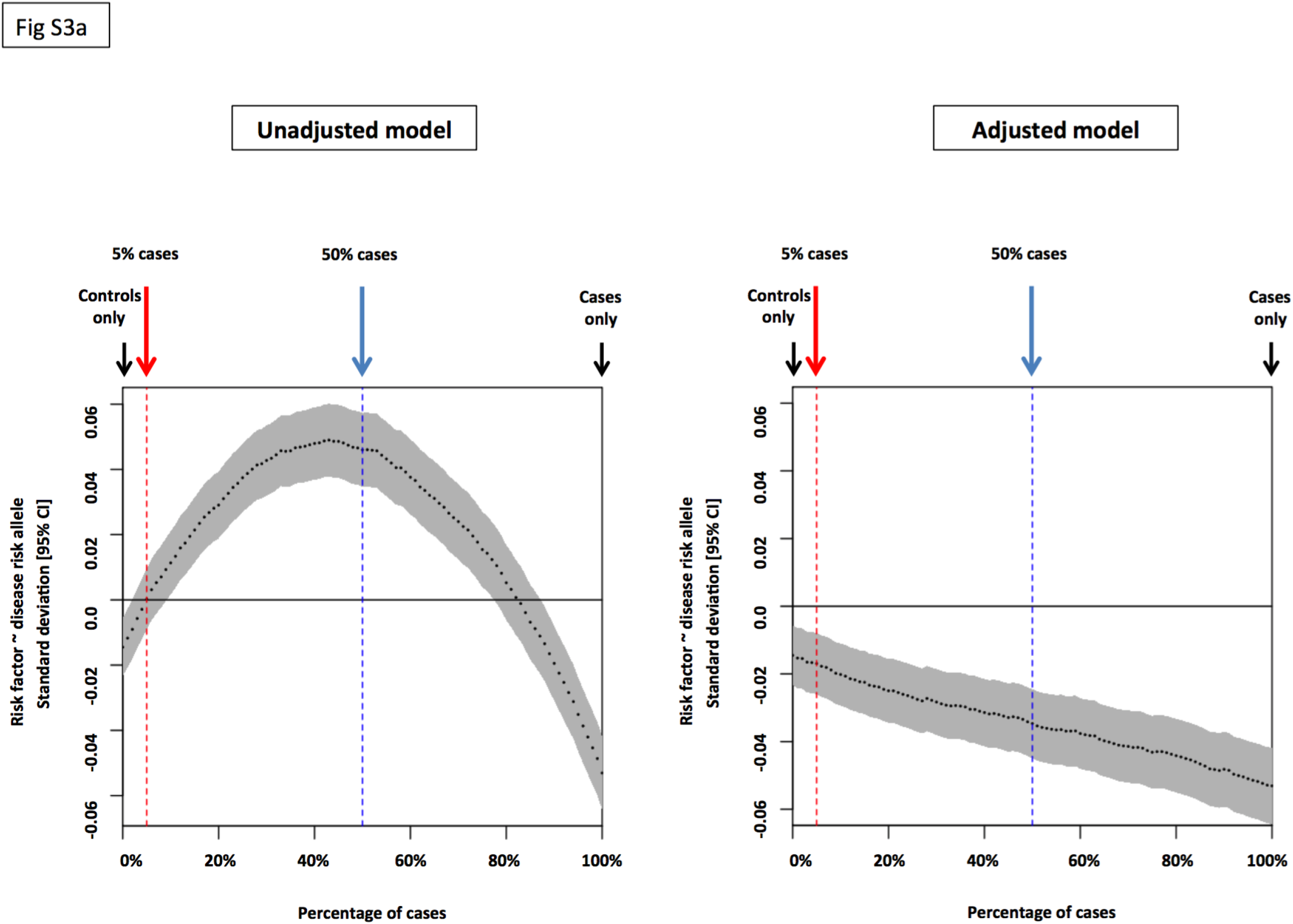
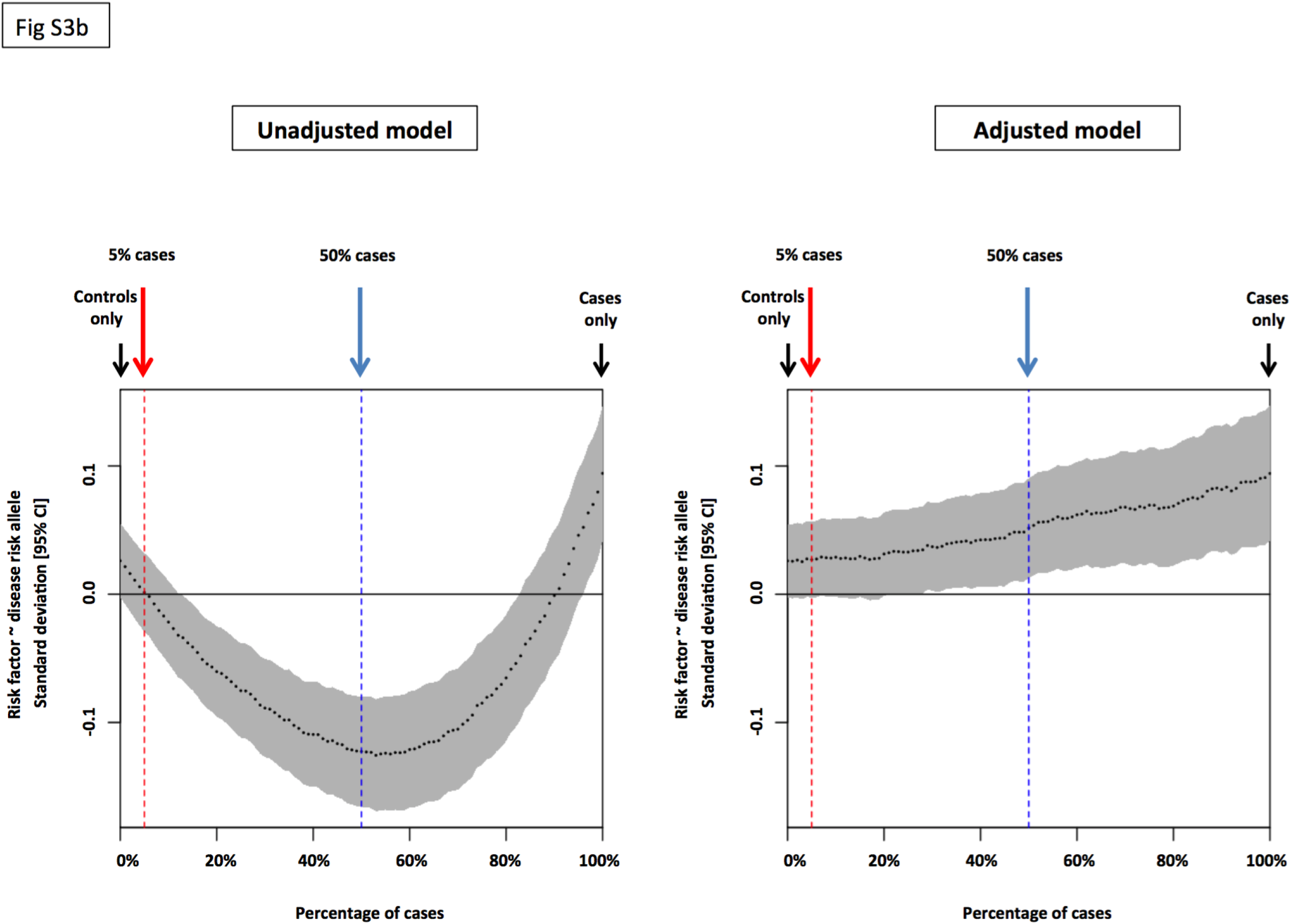
Enrichment for cases or controls produces spurious association. Enrichment for cases or controls produces spurious evidence of association between disease risk alleles and a risk factor correlated with the disease (2.5 OR per SD) in a scenario (**a**) where a risk allele (MAF 30%) increases risk with an odds ratio of 1.4 (similar to the *TCF7L2* type 2 diabetes scenario) and (**b**) where a protective allele (MAF 2%) reduces risk of disease with an odds ratio of 0.5 (similar to the *CCND2* type 2 diabetes scenario) in two models: unadjusted for disease status (left) and adjusted for disease status (right. The grey area represents 95% confidence interval (CI) around the effect estimate. The data is from simulated population of 100,000 individuals.

**Supplementary Figure 4.**
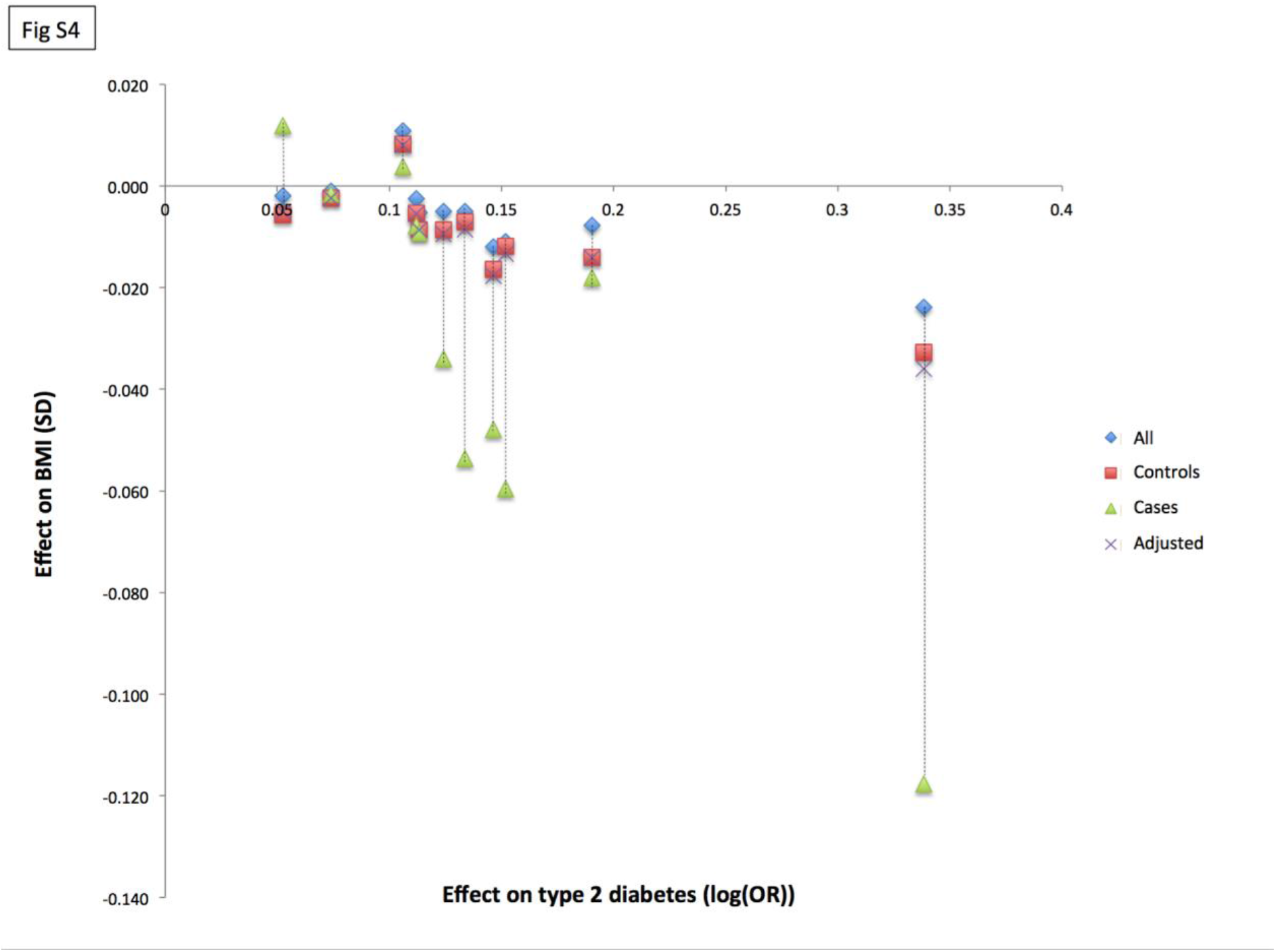
Real examples of index event bias. Examples of index event biased effect of the 11 insulin secretion SNPs on BMI (SD) in the UK Biobank study.

**Supplementary Figure 5.**
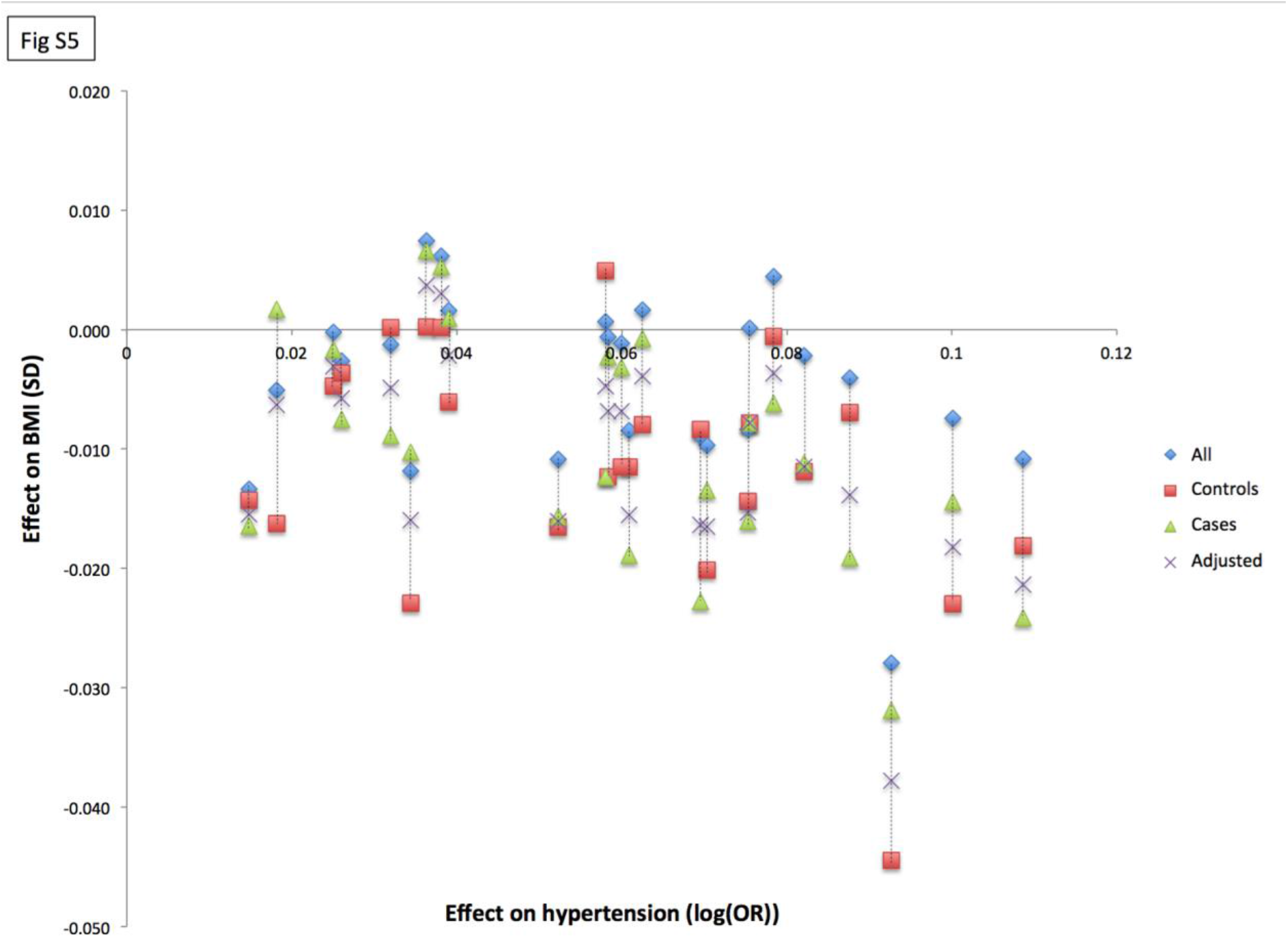
Real examples of index event bias. Examples of index event biased effect of the 25 blood pressure SNPs on BMI (SD) in the UK Biobank study.

**Supplementary Figure 6.**
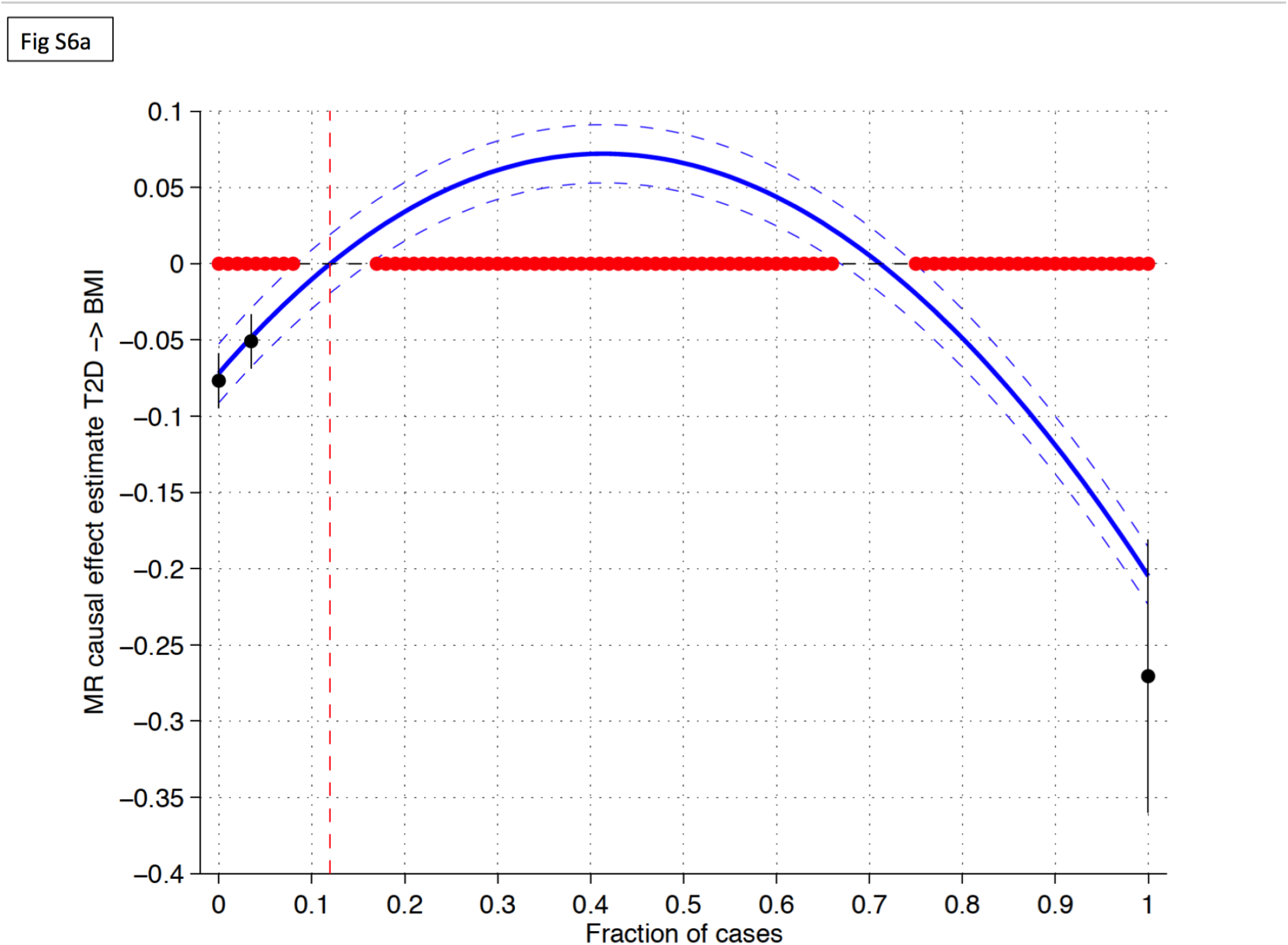
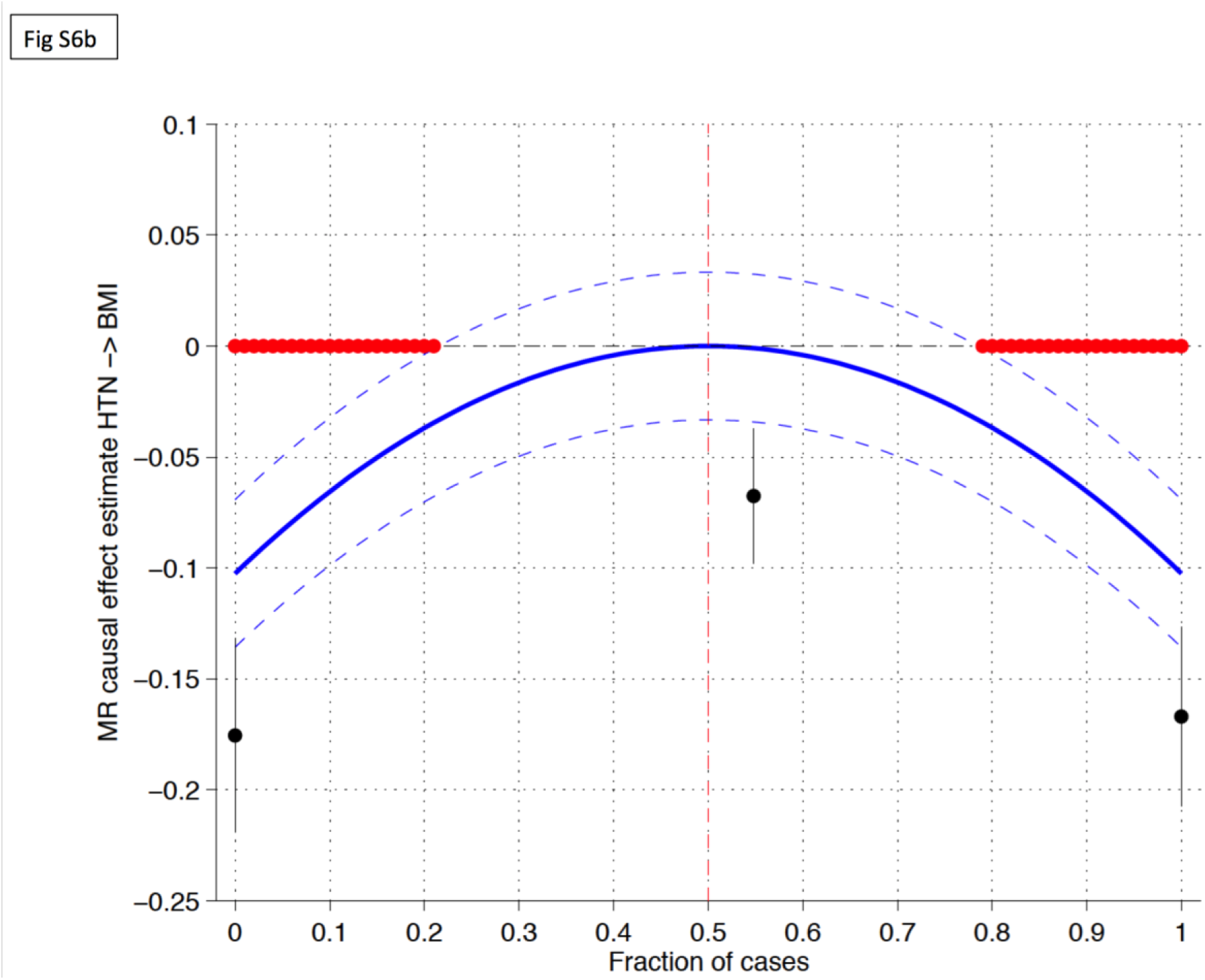
Index event bias in Mendelian randomization studies. **(a)** Type 2 diabetes (T2D) -> BMI causal effect estimates as a function of type 2 diabetes disease prevalence. We used the analytical formula for index event bias to perform Mendelian randomization (MR) using the 11 type 2 diabetes SNPs as instruments in order to derive estimate for the causal effect of type 2 diabetes on BMI based on parameters obtained from the UK Biobank data. The formula allowed causal effect estimation for the whole range of disease prevalences (0–100%). We overlaid the MR estimates obtained from the UK Biobank (all sample, controls only, cases only) and marked them with black disks. Estimates from all subsamples indicate significant negative causal effect. However, the comparison to the IEB curves reveals that the MR estimates from UK Biobank are compatible with pure index event bias with no causal effect. **(b)** Same analysis was done for hypertension (HTN) -> BMI causal effect estimates as a function of hypertension prevalence. This MR used 25 hypertension SNPs as instruments in order to derive estimate for the causal effect of hypertension on BMI. The comparison to the IEB curves reveals that the MR estimates from UK Biobank are not compatible with pure index event bias and a distinct negative causal effect is detectable.

## Supplementary Tables

**Supplementary Table 1.** The 11 SNPs associated with type 2 diabetes through insulin secretion^19^. *EFFECT_ALLELE = Impaired insulin secretion allele.

| SNP | LOCUS | CHR | POSITION | BUILD | EFFECT ALLELE* | OTHER ALLELE |
| --- | --- | --- | --- | --- | --- | --- |
| rs7903146 | *TCF7L2* | 10 | 114748339 | b36 | T | C |
| rs10965250 | *CDKN2A/B* | 9 | 22123284 | b36 | G | A |
| rs11899863 | *THADA* | 2 | 43472323 | b36 | C | T |
| rs10440833 | *CDKALI* | 6 | 20796100 | b36 | A | T |
| rs5015480 | *HHEX/IDE* | 10 | 94455539 | b36 | C | T |
| rs3802177 | *SLC30A8* | 8 | 118254206 | b36 | G | A |
| rs4607517 | *GCK* | 7 | 44202193 | b36 | A | G |
| rs2191349 | *DGKB/TMEM195* | 7 | 15030834 | b36 | T | G |
| rs10830963 | *MTNR1B* | 11 | 92348358 | b36 | G | C |
| rs340874 | *PROX1* | 1 | 212225879 | b36 | C | T |
| rs11708067 | *ADCY5* | 3 | 124548468 | b36 | A | G |

**Supplementary Table 2.**
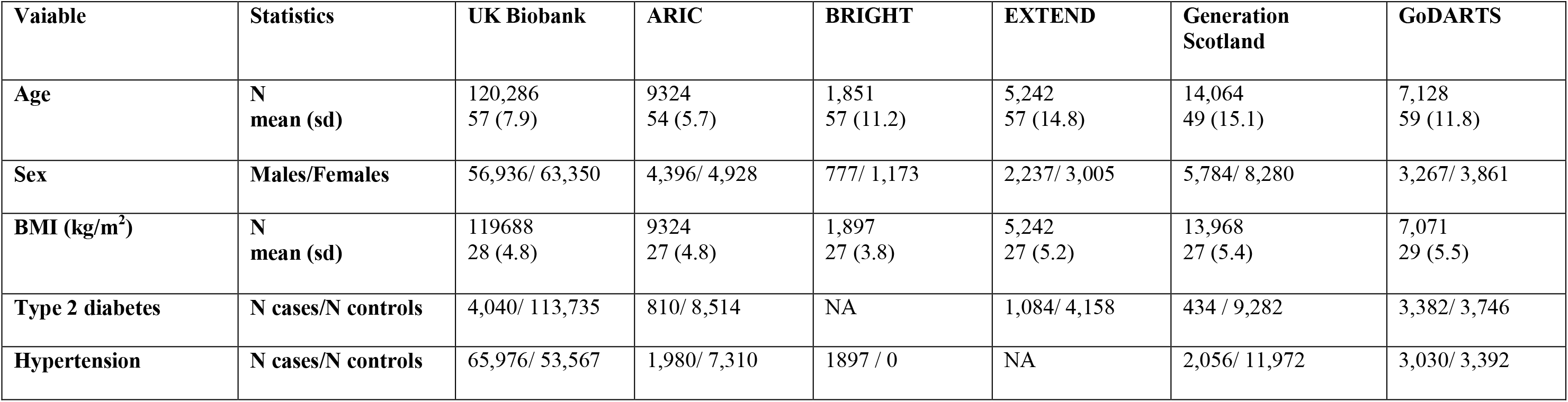
Summary details and relevant characteristics.

**Supplementary Table 3.** 25 SNPs associated with systolic blood pressure^25^. *EFFECT_ALLELE = Blood pressure increasing allele.

| SNP | LOCUS | CHR | POSITION | BUILD | EFFECT ALLELE* | OTHER ALLELE |
| --- | --- | --- | --- | --- | --- | --- |
| rs7129220 | *ADM* | 11 | 10,307,114 | 36 | A | G |
| rs17249754 | *ATP2B1* | 12 | 88,584,717 | 36 | G | A |
| rs805303 | *BAT2/BAT5* | 6 | 31,724,345 | 36 | G | A |
| rs4590817 | *C10orf107* | 10 | 63,137,559 | 36 | G | C |
| rs1813353 | *CACNB2(3')* | 10 | 18,747,454 | 36 | T | C |
| rs4373814 | *CACNB2(5')* | 10 | 18,459,978 | 36 | C | G |
| rs11191548 | *CYP17A1/NT5C2* | 10 | 104,836,168 | 36 | T | C |
| rs1378942 | *CYP11A1/ULK3* | 15 | 72,864,420 | 36 | C | A |
| rs11953630 | *EBF1* | 5 | 157,777,980 | 36 | C | T |
| rs1458038 | *FGF5* | 4 | 81,383,747 | 36 | T | C |
| rs633185 | *FLJ32810/TMEM133* | 11 | 100,098,748 | 36 | C | G |
| rs2521501 | *FURIN/FES* | 15 | 89,238,392 | 36 | T | A |
| rs6015450 | *GNAS/EDN3* | 20 | 57,184,512 | 36 | G | A |
| rs17608766 | *GOSR2* | 17 | 42,368,270 | 36 | C | T |
| rs1799945 | *HFE* | 6 | 26,199,158 | 36 | G | C |
| rs1327235 | *JAG1* | 20 | 10,917,030 | 36 | G | A |
| rs419076 | *MECOM* | 3 | 170,583,580 | 36 | T | C |
| rs2932538 | *MOV10* | 1 | 113,018,066 | 36 | G | A |
| rs17367504 | *MTHFR/NPPB* | 1 | 11,785,365 | 36 | A | G |
| rs1173771 | *NPR3/C5orf23* | 5 | 32,850,785 | 36 | G | A |
| rs932764 | *PLCE1* | 10 | 95,885,930 | 36 | G | A |
| rs381815 | *PLEKHA7* | 11 | 16,858,844 | 36 | T | C |
| rs3184504 | *SH2B3* | 12 | 110,368,991 | 36 | T | C |
| rs10850411 | *TBX5/TBX3* | 12 | 113,872,179 | 36 | T | C |
| rs12940887 | *ZNF652* | 17 | 44,757,806 | 36 | T | C |

**Supplementary Table 4.** The effect of 11 insulin secretion SNPs, BMI and the insulin secretion genetic risk score on risk of type 2 diabetes. OR: odds ration; LCI: lower confidence interval; UCI: upper confidence interval; P: p-value; N: total sample size.

|  | UK Biobank study |  |  |  |  | Additional studies |  |  |  |  | Meta-analyses of all studies |  |  |  |  |
| --- | --- | --- | --- | --- | --- | --- | --- | --- | --- | --- | --- | --- | --- | --- | --- |
| EXPOSURE | OR | LCI | UCI | P | N | OR | LCI | UCI | P | N | OR | LCI | UCI | P | N |
| *TCF7L2* | 1.40 | 1.34 | 1.47 | 3E-45 | 117688 | 1.36 | 1.29 | 1.43 | 7E-30 | 30725 | 1.38 | 1.34 | 1.43 | 1E-75 | 148413 |
| *THADA* | 1.16 | 1.08 | 1.26 | 2E-4 | 117688 | 1.19 | 1.09 | 1.30 | 0.0002 | 25668 | 1.17 | 1.11 | 1.24 | 7E-8 | 143356 |
| *CDKN2AB* | 1.21 | 1.14 | 1.29 | 3E-9 | 117688 | 1.13 | 1.05 | 1.20 | 0.0005 | 28940 | 1.17 | 1.12 | 1.23 | 6E-12 | 146628 |
| *SLC30A8* | 1.13 | 1.08 | 1.19 | 1E-8 | 117688 | 1.17 | 1.11 | 1.23 | 2E-8 | 28959 | 1.15 | 1.11 | 1.19 | 1E-14 | 146647 |
| *CDKALI* | 1.16 | 1.10 | 1.22 | 6E-9 | 117688 | 1.09 | 1.04 | 1.15 | 0.0008 | 30734 | 1.12 | 1.08 | 1.16 | 3E-10 | 148422 |
| *MTNR1B* | 1.11 | 1.06 | 1.17 | 2E-5 | 117688 | 1.10 | 1.04 | 1.16 | 0.0009 | 30466 | 1.11 | 1.07 | 1.15 | 8E-08 | 148154 |
| *HHEX* | 1.14 | 1.09 | 1.20 | 1E-8 | 117688 | 1.07 | 1.02 | 1.12 | 0.01 | 30738 | 1.10 | 1.07 | 1.14 | 9E-09 | 148426 |
| *GCK* | 1.12 | 1.06 | 1.19 | 9E-5 | 117688 | 1.09 | 1.01 | 1.16 | 0.02 | 25662 | 1.11 | 1.06 | 1.16 | 0.000006 | 143350 |
| *PROX1* | 1.08 | 1.03 | 1.13 | 0.001 | 117688 | 1.07 | 1.01 | 1.13 | 0.01 | 23883 | 1.08 | 1.04 | 1.12 | 0.00006 | 141571 |
| *ADCY5* | 1.05 | 1.00 | 1.11 | 0.05 | 117688 | 1.12 | 1.06 | 1.20 | 0.0001 | 25657 | 1.08 | 1.04 | 1.12 | 0.0002 | 143345 |
| *DGKB* | 1.12 | 1.07 | 1.17 | 1E-6 | 117688 | 0.97 | 0.92 | 1.02 | 0.2 | 21091 | 1.05 | 1.02 | 1.09 | 0.003 | 138779 |
| BMI (per SD) | 2.47 | 2.39 | 2.55 | 0 | 117120 | 2.30 | 2.21 | 2.38 | 0 | 34974 | 2.39 | 2.34 | 2.45 | 0 | 152094 |
| Insulin secretion<br>genetic risk score | 1.24 | 1.21 | 1.28 | 7E-44 | 117688 | 1.19 | 1.15 | 1.23 | 2E-23 | 30486 | 1.22 | 1.19 | 1.25 | 2E-72 | 148174 |

**Supplementary Table 5.**
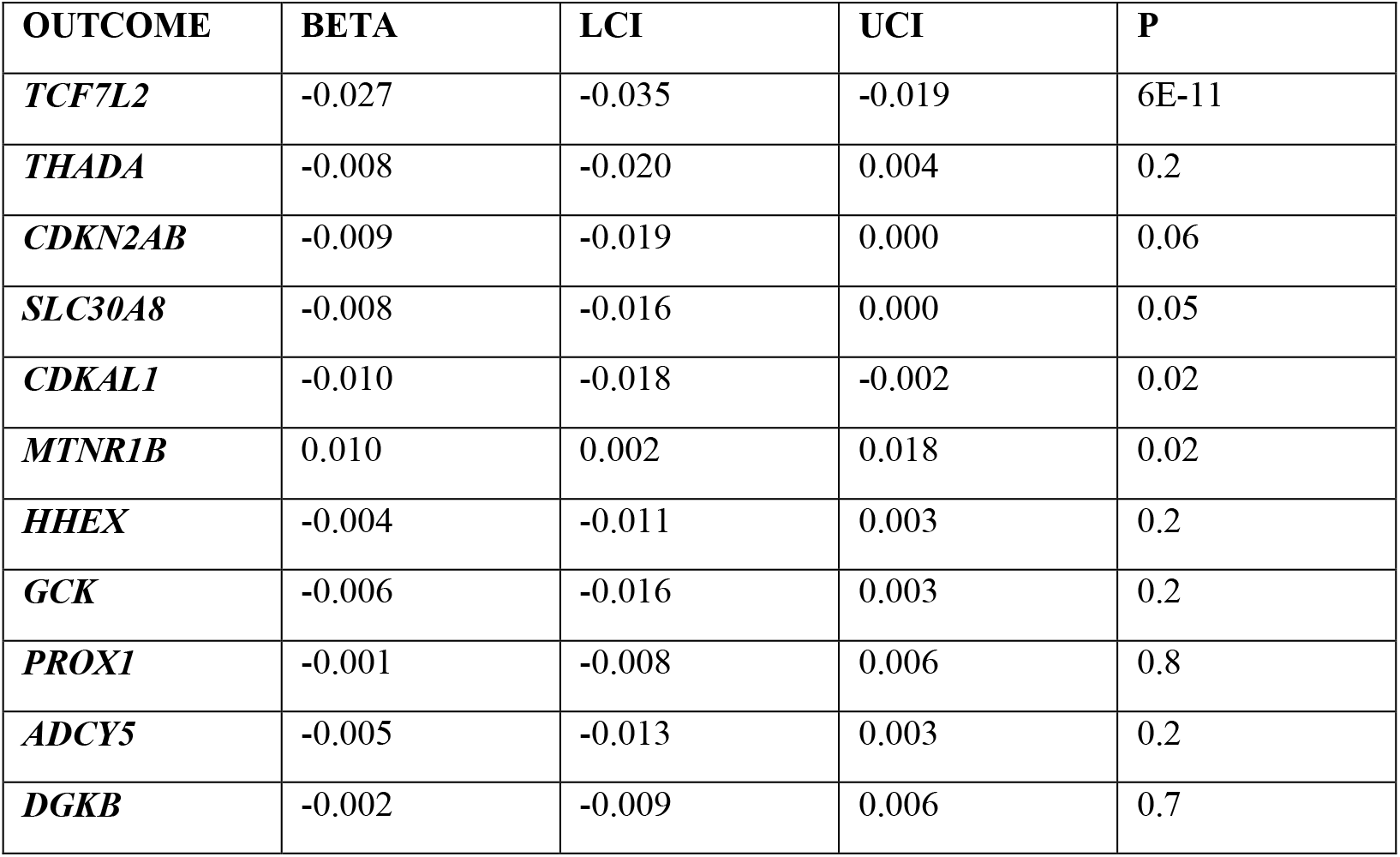
Association between 11 insulin secretion SNPs and BMI (SD). Result is from the meta-analysis of the UK Biobank (all individuals unstratified and unadjusted) and 4 additional studies: Generation Scotland (all individuals unstratified and unadjusted), GoDarts (cases and controls analysed as strata), EXTEND (cases and controls analysed as strata) and ARIC (all individuals unstratified and unadjusted). LCI: lower confidence interval; UCI: upper confidence interval; P: p-value.

| **OUTCOME** | **BETA** | **LCI** | **UCI** | **P** |
| --- | --- | --- | --- | --- |
| ***TCF7L2*** | -0.027 | -0.035 | -0.019 | 6E-11 |
| ***THADA*** | -0.008 | -0.020 | 0.004 | 0.2 |
| ***CDKN2AB*** | -0.009 | -0.019 | 0.000 | 0.06 |
| ***SLC30A8*** | -0.008 | -0.016 | 0.000 | 0.05 |
| ***CDKALI*** | -0.010 | -0.018 | -0.002 | 0.02 |
| ***MTNR1B*** | 0.010 | 0.002 | 0.018 | 0.02 |
| ***HHEX*** | -0.004 | -0.011 | 0.003 | 0.2 |
| ***GCK*** | -0.006 | -0.016 | 0.003 | 0.2 |
| ***PROX1*** | -0.001 | -0.008 | 0.006 | 0.8 |
| ***ADCY5*** | -0.005 | -0.013 | 0.003 | 0.2 |
| ***DGKB*** | -0.002 | -0.009 | 0.006 | 0.7 |

**Supplementary Table 6.** The effect of 25 blood pressure SNPs, BMI and the blood pressure genetic risk score on risk of hypertension. OR: odds ration; LCI: lower confidence interval; UCI: upper confidence interval; P: p-value; N: total sample size.

|  | UK Biobank study |  |  |  |  | Additional studies |  |  |  |  | Meta-analyses of all studies |  |  |  |  |
| --- | --- | --- | --- | --- | --- | --- | --- | --- | --- | --- | --- | --- | --- | --- | --- |
| EXPOSURE | OR | LCI | UCI | P | N | OR | LCI | UCI | P | N | OR | LCI | UCI | P | N |
| *FGF5* | 1.11 | 1.08 | 1.13 | 5E-25 | 119504 | 1.12 | 1.07 | 1.18 | 5E-07 | 29508 | 1.11 | 1.09 | 1.14 | 6E-25 | 149012 |
| *MTHFR* | 1.11 | 1.09 | 1.14 | 5E-20 | 119504 | 1.11 | 1.05 | 1.18 | 0.0002 | 29567 | 1.11 | 1.09 | 1.13 | 6E-23 | 149071 |
| *CYP17A1* | 1.10 | 1.06 | 1.13 | 2E-8 | 119504 | 1.14 | 1.06 | 1.23 | 0.0006 | 29579 | 1.11 | 1.07 | 1.14 | 2E-11 | 149083 |
| *GNAS* | 1.09 | 1.06 | 1.12 | 1E-10 | 119504 | 1.12 | 1.06 | 1.19 | 0.0002 | 29570 | 1.10 | 1.07 | 1.12 | 1 E-12 | 149074 |
| *ATP2B1* | 1.09 | 1.06 | 1.11 | 2E-12 | 119504 | 1.13 | 1.05 | 1.21 | 0.0008 | 23163 | 1.09 | 1.07 | 1.12 | 1E-15 | 142667 |
| *FURIN* | 1.08 | 1.06 | 1.10 | 2E-15 | 119504 | 1.08 | 1.02 | 1.13 | 0.003 | 25434 | 1.08 | 1.06 | 1.10 | 5E-18 | 144938 |
| *FLJ32810* | 1.08 | 1.06 | 1.10 | 1E-14 | 119504 | 1.10 | 1.05 | 1.15 | 0.0001 | 25243 | 1.08 | 1.06 | 1.10 | 2E-19 | 144747 |
| *NPR3* | 1.08 | 1.06 | 1.10 | 2E-18 | 119504 | 1.06 | 1.02 | 1.11 | 0.005 | 25532 | 1.08 | 1.06 | 1.10 | 9E-18 | 145036 |
| *CACNB2_3* | 1.07 | 1.05 | 1.09 | 6E-14 | 119504 | 1.06 | 1.01 | 1.12 | 0.03 | 23162 | 1.07 | 1.05 | 1.09 | 1E-13 | 142666 |
| *SH2B3* | 1.07 | 1.05 | 1.09 | 3E-15 | 119504 | 1.00 | 0.95 | 1.05 | 0.95 | 19118 | 1.06 | 1.04 | 1.08 | 2E-11 | 138622 |
| *GOSR2* | 1.06 | 1.04 | 1.09 | 1E-6 | 119504 | 1.10 | 1.03 | 1.16 | 0.002 | 29578 | 1.06 | 1.04 | 1.09 | 1E-8 | 149082 |
| *ADM* | 1.06 | 1.03 | 1.09 | 2E-5 | 119504 | 1.05 | 0.98 | 1.12 | 0.1 | 29512 | 1.06 | 1.03 | 1.09 | 0.00002 | 149016 |
| *EBF1* | 1.06 | 1.04 | 1.08 | 2E-10 | 119504 | 1.06 | 1.01 | 1.11 | 0.01 | 25460 | 1.06 | 1.04 | 1.08 | 5E-11 | 144964 |
| *HFE* | 1.06 | 1.04 | 1.09 | 4E-7 | 119504 | 1.07 | 1.01 | 1.13 | 0.02 | 29486 | 1.06 | 1.04 | 1.08 | 9E-8 | 148990 |
| *CYP11A1* | 1.05 | 1.03 | 1.07 | 3E-8 | 119504 | 1.09 | 1.04 | 1.14 | 0.0001 | 29577 | 1.06 | 1.04 | 1.07 | 9E-10 | 149081 |
| *C10orf107* | 1.06 | 1.04 | 1.09 | 2E-7 | 119504 | 1.00 | 0.94 | 1.05 | 0.9 | 29463 | 1.05 | 1.03 | 1.07 | 9E-6 | 148967 |
| **MECOM** | 1.04 | 1.02 | 1.06 | 1E-5 | 119504 | 1.06 | 1.01 | 1.11 | 0.01 | 23162 | 1.04 | 1.02 | 1.06 | 4E-6 | 142666 |
| **MOV10** | 1.04 | 1.02 | 1.06 | 3E-4 | 119504 | 1.05 | 1.00 | 1.10 | 0.06 | 29575 | 1.04 | 1.02 | 1.06 | 0.00001 | 149079 |
| **ZNF652** | 1.04 | 1.02 | 1.06 | 3E-5 | 119504 | 1.03 | 0.99 | 1.07 | 0.2 | 29577 | 1.04 | 1.02 | 1.06 | 0.00003 | 149081 |
| **CACNB2_5** | 1.03 | 1.02 | 1.05 | 1E-4 | 119504 | 1.05 | 1.00 | 1.11 | 0.05 | 23106 | 1.03 | 1.02 | 1.05 | 0.00001 | 142610 |
| **TBX5** | 1.03 | 1.01 | 1.05 | 0.006 | 119504 | 1.02 | 0.97 | 1.06 | 0.5 | 29516 | 1.03 | 1.01 | 1.05 | 0.003 | 149020 |
| **PLCE1** | 1.03 | 1.01 | 1.05 | 3E-4 | 119504 | 1.01 | 0.97 | 1.05 | 0.6 | 29519 | 1.03 | 1.01 | 1.04 | 0.004 | 149023 |
| **JAG1** | 1.03 | 1.01 | 1.04 | 0.005 | 119504 | 1.02 | 0.97 | 1.06 | 0.5 | 25532 | 1.03 | 1.01 | 1.04 | 0.00007 | 145036 |
| **PLEKHA7** | 1.02 | 1.00 | 1.04 | 0.06 | 119504 | 1.04 | 0.98 | 1.10 | 0.2 | 23161 | 1.02 | 1.00 | 1.04 | 0.02 | 142665 |
| **BAT2** | 1.01 | 1.00 | 1.03 | 0.1 | 119504 | 1.02 | 0.98 | 1.06 | 0.3 | 29563 | 1.01 | 1.00 | 1.03 | 0.1 | 149067 |
| **BMI (per SD)** | 1.63 | 1.61 | 1.65 | 0 | 118923 | 1.57 | 1.52 | 1.62 | 2E-190 | 29575 | 1.62 | 1.60 | 1.64 | <E-200 | 148498 |
| **Hypertension**<br>**genetic risk**<br>**score** | 1.20 | 1.18 | 1.21 | 8E-180 | 119504 | 1.20 | 1.16 | 1.24 | 7E-31 | 25053 | 1.20 | 1.19 | 1.21 | 6E-207 | 144557 |

**Supplementary Table 7.** Association between 25 blood pressure SNPs and BMI (SD). Result is from the meta-analysis of the UK Biobank (all individuals unstratified and unadjusted) and 4 additional studies: Generation Scotland (all individuals unstratified and unadjusted) EXTEND (all individuals unstratified and unadjusted), BRIGHT (cases only) and ARIC (all individuals unstratified and unadjusted). LCI: lower confidence interval; UCI: upper confidence interval; P: p-value.

| OUTCOME | BETA | LCI | UCI | P |
| --- | --- | --- | --- | --- |
| *FGF5* | -0.002 | -0.010 | 0.006 | 0.6 |
| *MTHFR* | -0.008 | -0.018 | 0.002 | 0.1 |
| *CYP17A1* | -0.025 | -0.039 | -0.012 | 0.0003 |
| *GNAS* | -0.004 | -0.015 | 0.007 | 0.5 |
| *ATP2B1* | -0.003 | -0.013 | 0.007 | 0.5 |
| *FURIN* | -0.002 | -0.010 | 0.006 | 0.6 |
| *FLJ32810* | -0.009 | -0.017 | 0.000 | 0.04 |
| *NPR3* | 0.004 | -0.004 | 0.012 | 0.3 |
| *CACNB2_3* | -0.009 | -0.017 | -0.001 | 0.04 |
| *SH2B3* | -0.010 | -0.017 | -0.002 | 0.01 |
| *GOSR2* | -0.002 | -0.013 | 0.008 | 0.7 |
| *ADM* | -0.002 | -0.014 | 0.009 | 0.7 |
| *EBF1* | -0.001 | -0.009 | 0.006 | 0.7 |
| *HFE* | 0.001 | -0.009 | 0.011 | 0.9 |
| *CYP1A1* | -0.012 | -0.020 | -0.004 | 0.002 |
| *C10orf107* | -0.006 | -0.016 | 0.004 | 0.2 |
| *MECOM* | 0.001 | -0.007 | 0.008 | 0.8 |
| *MOV10* | 0.008 | 0.000 | 0.016 | 0.06 |
| *ZNF652* | 0.009 | 0.001 | 0.016 | 0.02 |
| *CACNB2_5* | -0.010 | -0.017 | -0.002 | 0.01 |
| *TBX5* | -0.004 | -0.012 | 0.004 | 0.4 |
| *PLCE1* | -0.003 | -0.011 | 0.004 | 0.4 |
| *JAG1* | -0.002 | -0.009 | 0.006 | 0.6 |
| *PLEKHA7* | -0.005 | -0.014 | 0.003 | 0.2 |
| *BAT2* | -0.012 | -0.020 | -0.005 | 0.002 |

